# MouseCircuits.org: An online repository to guide the circuit era of disordered affect

**DOI:** 10.1101/2020.02.16.951608

**Authors:** Kristin R. Anderson, Dani Dumitriu

## Abstract

Affective disorders rank amongst the most disruptive and prevalent psychiatric diseases, resulting in enormous societal and economic burden, and immeasurable personal costs. Novel therapies are urgently needed but have remained elusive. The era of circuit-mapping in rodent models of disordered affect, ushered in by recent technological advancements allowing for precise and specific neural control, has reenergized the hope for precision psychiatry. Here, we present a novel whole-brain cumulative network and critically access the progress made to-date on circuits mediating affective-like behaviors in rodents to seek unifying principles of this cumulative data. We identified 106 original manuscripts in which optogenetics or chemogenetics were used to dissect behaviors related to fear-like, depressive-like or anxiety-like behaviors in rodents. Focusing on the 60 manuscripts that investigated pathways rather than regions, we identified emergent themes. We found that while a few pathways have been validated across similar behaviors and multiple labs, the data is mostly disjointed, with evidence of bidirectional effects of several pathways. Additionally, there is a need for analysis informed by observation prior to perturbation. Given the complex nature of brain connectivity, we argue that the compartmentalized viewpoint that develops as a consequence of fragmented pathway-specific manipulations does not readily lend itself to an integrative picture. To address this, we launched an interactive online consortium, MouseCircuits.org, an open-source platform for consolidated circuit data. This tool aims to support the shared vision of informed circuit dissection that ultimately leads to prevention and treatment of human disorders.

## The ERA OF CIRCUIT MAPPING

Mood disorders account for most years lost to disability (1). There is an urgent need for new effective therapeutics, but translation of laboratory discoveries to treatments for human disorders has thus far proven difficult. Excitingly, we are on the brink of a historical turning point. Until recently – borrowing from groundbreaking advancement of other organ systems – mechanistic dissection of disordered affect has targeted singular neurotransmitter systems, brain regions, or genes. This approach enabled the understanding of the individual building blocks of brain function and continues to be supported by novel theories involving global mechanism of affect disruption, implicating for example the immune system and the gut microbiome (2,3). However, the mysteries of the brain, a structure with idiosyncratic and interconnected architecture, are unlikely to be revealed solely on the basis of this type of sledgehammer approach.

### Enter the era of circuit dissection

In the last decade, groundbreaking technological advances have allowed neuroscientists to take control of neural firing with impressive precision and specificity **(Figure 1)**(4–7). An in-depth description of these tools is outside the scope of this paper [see (4–8) for comprehensive descriptions]. Concisely, two techniques have fundamentally changed the landscape of neural circuit dissection: optogenetics, which controls neuronal firing with light, and chemogenetics, which alters neuronal firing with otherwise biologically inert compounds. Optogenetics uses genetically engineered transmembrane channels that open in response to specific wavelengths of light to allow selective passage of charged ions to either depolarize or hyperpolarize targeted neurons. Chemogenetics uses G-protein coupled receptors known as Designer Receptors Exclusively Activated by Designer Drugs (DREADDs) to enhance or inhibit neuronal excitability in the presence of biologically inert compounds, such as clozapine N-oxide (CNO). Both tools can selectively activate or inactivate neurons of interest when combined with genetic labeling techniques. Optogenetic control provides high temporal precision on scales relevant to manipulations of individual action potentials. Chemogenetic control acts on slower timescales (minutes, hours, or days) but has the advantage of modulating endogenous activity by placing the neuron in either a depolarized state, for increased excitability, or hyperpolarized state, for decreased excitability.

**Fig. 1.**
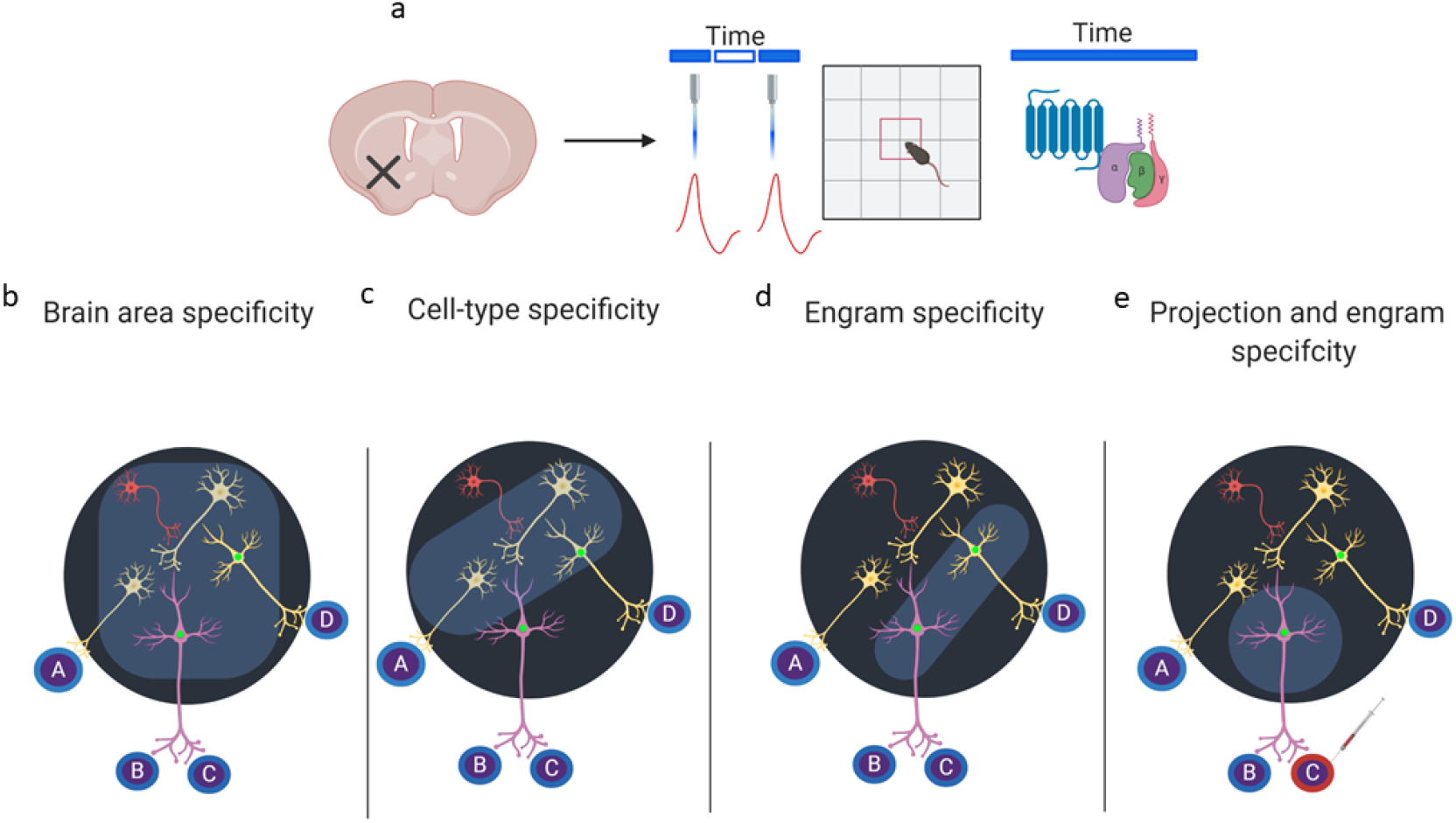
Evolution of the era of circuit mapping: from regional mapping to increasingly refined circuit dissection. (a) The neural basis of behavior was initially determined by non-specific lesions followed by the era of circuit mapping with optogenetic and chemogenetic manipulations. Optogenetics and chemogenetics have several key differences, with timing being the most relevant to behavioral manipulations. Optogenetics can be regulated on a second timescale while chemogenetics relies on Gprotein receptor dynamics. (b) Regional specificity of optogenetic and chemogenetic manipulations can be achieved with careful viral injections into a brain region of interest (light blue shading). (c) Combining optogenetics/chemogenetics with genetic targeting tools allows for cell-type specificity within regions of interest (light blue shading). (d) Growing use of activity-dependent targeting (green nuclei) further increased specificity of manipulations to temporally defined cell populations within a region. (e) Novel approaches using multiple systems such as Cre/loxp, inducible promoters, retrograde/intersectional/transsynaptic labeling allow for additional layers of specificity that combine cell-type, projection-type and/or temporally-defined targeting of neuronal populations.

Initial efforts used chemogenetic and optogenetic control of brain regions and confirmed findings from classical lesion studies (ex: (9)). Rapid progress followed, with increasingly refined targeting of neurons based on transcriptional-, projection- and/or activity-specificity **(Figure 1)**. However, despite substantial amassed data, knowledge of individual pathways exists largely in a realm that ignores the complexity of an interconnected whole-brain network. Therefore, at the moment, it is difficult to envision how these findings can be translated to improve the prevention and treatment of human mental health conditions. The goal of this paper is therefore threefold: (1) Summarize the cumulative knowledge of affective circuits obtained using chemogenetic and optogenetic manipulations in animal models; (2) Critically assess the current state of the era of circuit mapping, its promise and its perils; and (3) Introduce an online platform to help consolidate and integrate rapidly growing neurocircuit data moving forward: MouseCircuits.org.

### Neural map criteria

Our goal was to develop a functional neural map of affective state circuity data, enabled by chemogenetics and optogenetics. We searched PubMed (accessed on November 20th, 2019) for combinations of keywords referencing circuit dissection tools (e.g. “chemogenetics”, “optogenetics”, “DREADDs”) and keywords referencing affect (e.g. “fear”, “anxiety”, “depression”) or specific behavioral tests commonly used to asses affective-like responses in rodents (e.g. “social defeat stress”, “elevated plus maze”, “tail suspension”). Studies on mouse and rats only were included. These studies covered the investigation of circuits spanning 33 regions **(Table 1)**. No date restrictions were used, but consistent with chemogenetic and optogenetic tools’ recent development, all 102 identified manuscripts were from 2010 or later **(Table 2)**. An additional three manuscripts were identified based on Twitter updates of BioRxiv preprints, for a total of 106 reviewed studies **(Table 2**, page 18).

**Table 1.** Region names and abbreviations from the Allen Brain Atlas with the expectation of the infralmibic cortex (IL rather than ILA) and the addition of the amygdala (Amyg), dorsal hippocampus (dHPC), ventral hippocampus (vHPC), and medial prefrontal cortex (mPFC).

| Abbreviation | Region |
| --- | --- |
| Amyg | Amygdala |
| ACC | Anterior cingulate cortex |
| Acx | Auditor cortex |
| BLA | Basolateral amygdala |
| BST | Bed nucleus of the stria terminalis |
| CeA | Central amygdala |
| DG | Dentate gyrus |
| DR | Dorsal raphe nucleus |
| EC | External capsule |
| H | Hypothalamus |
| HPF | Hippocampus formation |
| dHPC | Dorsal Hippocampus |
| vHPC | Ventral Hippocampus |
| AI | Insula |
| IL | Infralimbic cortex |
| ILT | Intralaminar thalamus |
| LA | Lateral amygdala |
| LC | Locus coeruleus |
| LDT | Laterodorsal tegmentum |
| LH | Lateral habenula |
| LS | Lateral septum |
| MDT | Mediodorsal nucleus of the thalamus |
| mPFC | Medial prefrontal cortex |
| ACB | Nucleus accumbens |
| NR | Nucleus reuniens |
| PAG | Periaqueductal gray |
| PL | Prelimbic cortex |
| PVT | Dorsal midline thalamus |
| PC | Piriform cortex |
| SC | Superior colliculus |
| T | Thalamus (including MFD) |
| VTA | Ventral tegmental area |
| VS | Ventral striatum |

**Table 2.** Studies of perturbed affect reviewed. Summary of studies reviewed and number assigned in Figure 3. Studies are presented in chronological order by publication date. See tables 3 and 4 for details. *Note: These numbers match* Figure 3, *not the above bibliography.*

| Figure 3 Number | Reference |
| --- | --- |
| 1 | Covington et al., 2010 |
| 2 | Johansen et al., 2010 |
| 3 | Tye et al., 2011 |
| 4 | Goshen et al., 2011 |
| 5 | Chaudhury et al.,2012 |
| Continued on next page |  |

| **Figure 3 Number** | **Reference** |
| --- | --- |
| 6 | Liu et al.,2012 |
| 7 | Warden et al.,2012 |
| 8 | Tye et al.,2013 |
| 9 | Xu and Südhof 2013 |
| 10 | Kim et al.,2013 |
| 11 | Jennings et al.,2013 |
| 12 | Felix-Ortiz et al.,2013 |
| 13 | Challis et al.,2013 |
| 14 | Lobo et al.,2013 |
| 15 | Felix-Ortiz and Tye 2014 |
| 16 | Senn et al.,2014 |
| 17 | Anthony et al.,2014 |
| 18 | Challis et al.,2014 |
| 19 | Zhu et al.,2014 |
| 20 | Friedman et al.,2014 |
| 21 | Sparta et al.,2014 |
| 22 | Ohmura et al.,2014 |
| 23 | Wolff et al.,2014 |
| 24 | Redondo et al.,2014 |
| 25 | Soumier and Sibile,2014 |
| 26 | Kwon et al.,2014 |
| 27 | Yizhar et al.,2014 |
| 28 | Bagot et al.,2015 |
| 29 | Christoffel et al.,2015 |
| 30 | Chase Francis et al.,2015 |
| 31 | Perova et al.,2015 |
| 32 | Sachs et al.,2015 |
| 33 | Do-Monte et al., 2015a |
| 34 | Do-Monte et al.,2015b |
| 35 | Penzo et al.,2015 |
| 36 | Ramirez et al.,2015 |
| 37 | Gore et al.,2015 |
| 38 | Namburi et al.,2015 |
| 39 | Carreno et al.,2015 |
| 40 | Teissier et al.,2015 |
| 41 | Yang et al.,2016a |
| 42 | Marcinkiewicz et al.,2016 |
| 43 | Yang et al.,2016b |
| 44 | Felix-Ortiz et al.,2016 |
| 45 | Rashid et al.,2016 |
| 46 | Kim et al.,2016a |
| 47 | Dejean et al., 2016 |
| 48 | Johnson et al.,2016 |
| 49 | Kim et al.,2016b |
| 50 | Wook Koo et al.,2016 |
| 51 | Urban et al.,2016 |
| 52 | Xu et al.,2016 |
| 53 | Zou et al.,2016 |
| 54 | Fadok et al.,2017 |
| 55 | Klavir et al.,2017 |
| 56 | Parfitt et al.,2017 |
| 57 | Vetere et al.,2017 |
| Continued on next page |  |

| **Figure 3 Number** | **Reference** |
| --- | --- |
| 58 | Asok et al.,2017 |
| 59 | Franklin et al.,2017 |
| 60 | Knowland et al.,2017 |
| 61 | McCall et al.,2017 |
| 62 | Arico et al.,2017 |
| 63 | Miller et al.,2017 |
| 64 | Assareh et al.,2017 |
| 65 | Dolzani et al.,2018 |
| 66 | Marek et al., 2018a |
| 67 | Jimenez et al.,2018 |
| 68 | Bloodgood et al.,2018 |
| 69 | DeNardo et al.,2018 |
| 70 | Marek et al., 2018b |
| 71 | Diehl et al.,2018 |
| 72 | Zhang et al.,2018 |
| 73 | Mukherjee and Caroni 2018 |
| 74 | Lowery-Gionta et al., 2018a |
| 75 | Lowery-Gionta et al.,2018b |
| 76 | Yamauchi et al.,2018 |
| 77 | Tipps et al.,2018 |
| 78 | Fernandez et al.,2018 |
| 79 | Anacker et al., 2018 |
| 80 | Salinas-Hernandez et al.,2018 |
| 81 | Bian et al.,2019 |
| 82 | Besnard et al.,2019 |
| 83 | Chen and Bi 2019 |
| 84 | Gehrlach et al.,2019 |
| 85 | Kato et al.,2019 |
| 86 | Vasquez et al.,2019 |
| 87 | Wang et al.,2019 |
| 88 | Ortiz et al.,2019 |
| 89 | Berg et al.,2019 |
| 90 | Bernard et al.,2019 |
| 91 | Rozeske et al.,2018 |
| 92 | Lacagnina et al.,2019 |
| 93 | Moda-Sava et al., 2019 |
| 94 | Zhang et al.,2019 |
| 95 | Matos et al.,2019 |
| 96 | Wilmot et al.,2019 |
| 97 | Roy et al.,2019 |
| 98 | Zhang et al.,2019b |
| 99 | Giannotti et al.,2019 |
| 100 | Sengupta et al.,2019 |
| 101 | Zhou et al.,2019 |
| 102 | Gutzeit et al.,2019 |
| 103 | Hartley et al.,2019 |
| 104 | Salvi et al.,2019 |
| 105 | Rogers-Carter et al.,2019 |
| 106 | Padilla-Coreano et al.,2019 |

## DISSECTION OF RODENT CIRCUITS FOR DISORDERED AFFECT

### Inclusion criteria for neural manipulations

A variety of tools can be used to manipulate neural activity including electric stimulation and transcranial magnetic stimulation. For the purpose of the creation of this neural network and guide, we restricted our search to studies in which optogenetics and/or chemogenetics were used to manipulate neuronal function, as these tools, when combined with genetic labeling, offer unprecedented specificity and control of microcircuits.

### Inclusion criteria for rodent behaviors

The relevance and success of rodent models for studying human mental health diseases is currently fiercely debated (2,10–18). It is well-established that animal models cannot recapitulate the heterogeneous and complex symptomology of patients with major depressive disorder (MDD), generalized anxiety disorder (GAD), post-traumatic stress disorder (PTSD), or any other disease of mood dysregulation. However, there is also general agreement that rodent models, when used appropriately, will continue to be a crucial intermediary step as we move treatments through the translational pipeline (17).

In the collection of data point for a whole-brain neural network we included rodent studies that share the broad conceptual framework of modeling core aspects of human affect. For feasibility, we did not search an exhaustive of all affective-like behaviors (e.g. “motivation” and “reward” were not included) but targeted three types of behavioral responses in rodents: “fear-like” (innate or learned), “anxiety-like”, and “depressive-like” **(Figure 2)**. These behaviors and – the affective states they are associated with – are not necessarily separable in either humans or animal models. There is a high degree of overlapping symptomology in different diseases, as well as comorbidity among disorders (19–22). Nevertheless, understanding the precise pathways involved in specific symptoms can generate important ground truth about affective circuits and supports the establishment of precision psychiatry. Recent introduction of Research Domain Criteria (RDoC) is a notable effort to move the field away from classification into disorders that reflect constellations of symptoms, such as MDD and GAD, and toward dimensions of functioning (23–26). It is hypothesized that constructs such as “acute threat (‘fear’)” and “potential threat (‘anxiety’)” share genetic, environmental, developmental, and neurocircuit eti ology across both diseases and species. Therefore, establishing a bird’s eye view of the various pathways with proven functional relevance to “acute threat” and “potential threat” in animal models is of crucial importance to the overarching vision of the RDoC framework.

**Fig. 2.**
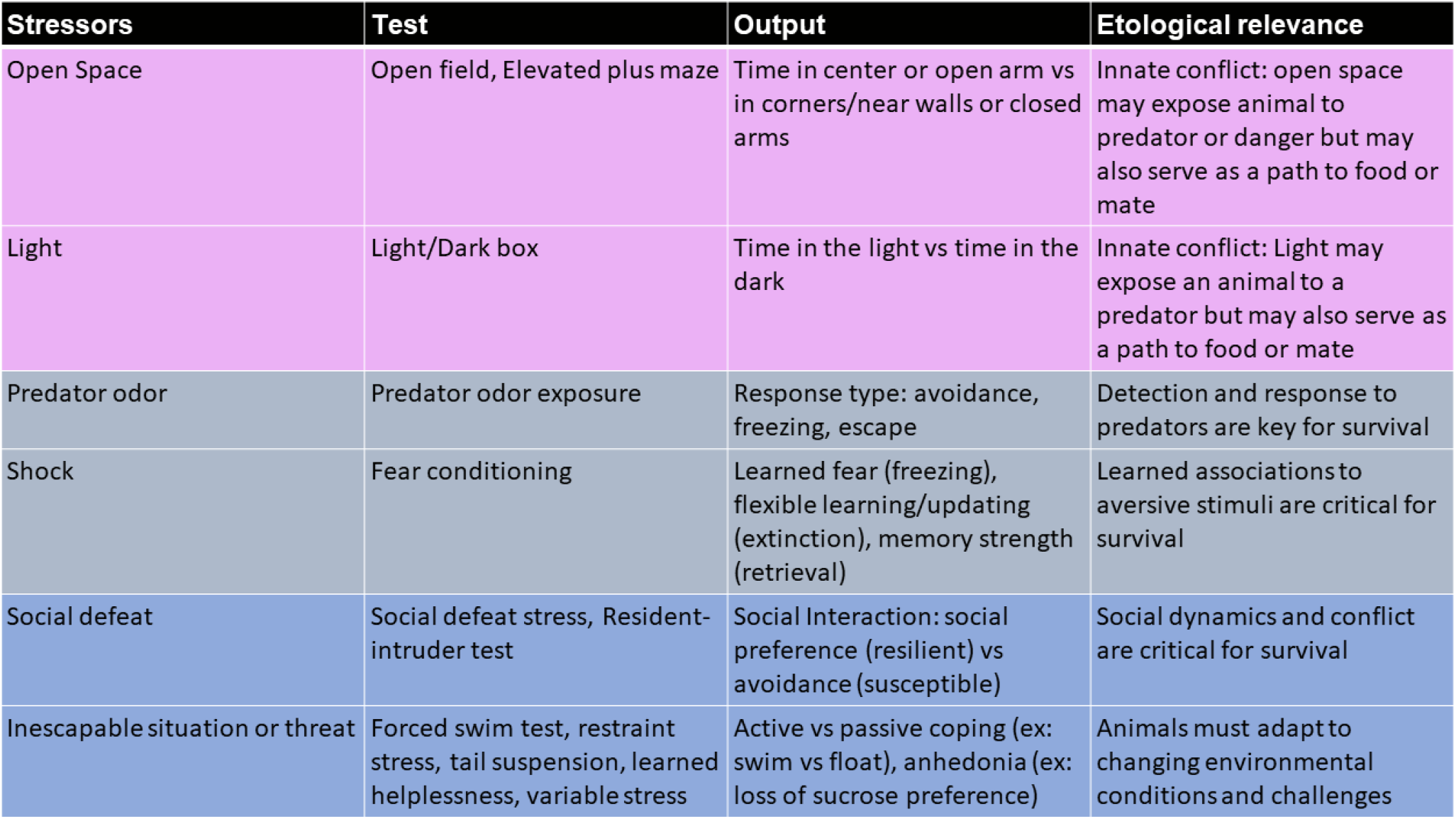
The behavioral toolbox. Summary of common behavioral models for affective-like states in rodent models. Purple rows are associated with anxiety-like behavior, light-blue rows are associated with fear-like behavior, and dark-blue rows are associated with depressive-like behaviors.

**Figure 3** shows the richness of rodent affective circuit data amassed to date. An interactive, searchable and updatable version of this figure will soon be available on MouseCircuits.org. Several themes emerge in the context of visualizing the cumulative investigated connectome of rodent affective-like behaviors.

**Fig. 3.**
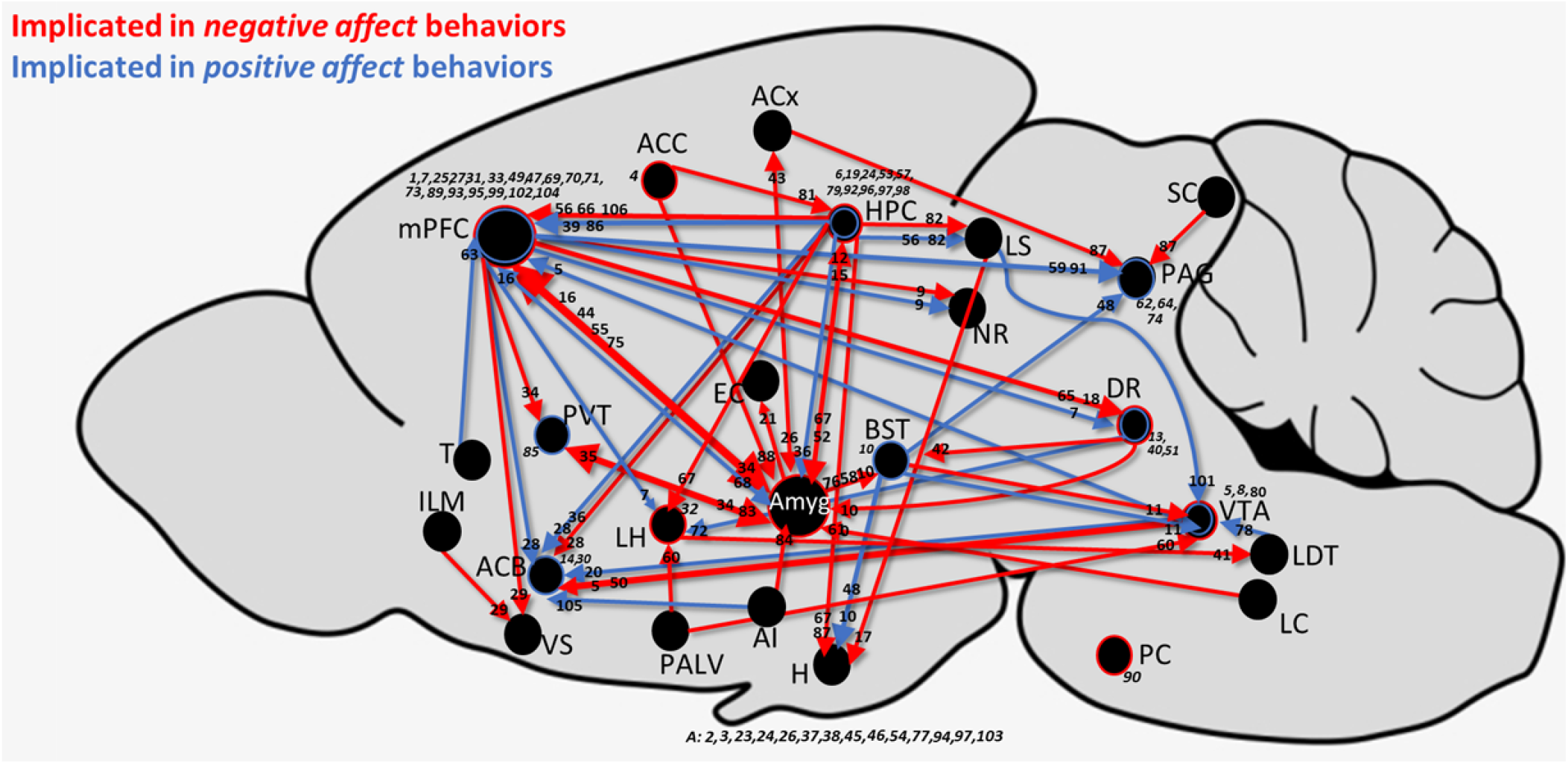
Verified rodent brain functional connectivity for affective-like behaviors. Pathway and region manipulations from 106 identified studies in which optogenetics and/or chemogenetics was used to probe a brain region’s or pathway’s contribution to anxiety-, fear-, or depressive-like behaviors, presented in the backdrop of the whole rodent brain. Red indicates regions and pathways in which activation promotes negative affective-like behavior, while blue indicates regions and pathways in which activation promotes positive affective-like behavior. Size of arrow corresponds to number of studies that have targeted a particular pathway.

### Landscape of studies dissecting rodent affective circuits

We identified a total of 106 studies investigating the role of brain regions or pathways in fear-like, anxiety-like and/or depressive-like behaviors using optogenetics or chemogenetics. Of these, 84% were mouse studies and 16% were rat studies. Optogenetics is more commonly used than chemogenetics, with region data investigated by optogenetics in 60% of studies, chemogenetics in 32% of studies, and both in 8% of studies. Pathway data was investigated by optogenetics in 70% of studies, chemogenetics in 20%, and both in 10% of studies. In both region and pathway investigations, the studies that utilized both methods (seven pathway focused and five region focused) obtained congruent results. Bidirectional control of the region (4 studies) or pathway of interest (13 studies) was investigated in 16% of the studies. Interestingly, only 6 of these studies (one region focused and five pathway focused) used the same method for bidirectional control. The remaining used optogenetics for activation and chemogenetics for inhibition or vice versa (three region focused and two pathway focused).

Males rodents are overwhelmingly more commonly studied, with 81% of studies using only males (35/49 region focused and 51/60 pathway focused). In the remaining studies, none used only females, 17 (9 region focused and 8 pathway focused) studies used both males and females, and 7 studies did not report sex (5 region focused and 2 pathway focused).

A total of 49 studies targeted 15 different brain regions **(Table 3)**. A minority of these regions have been looked at in the context of multiple types of affective behaviors. The three most commonly studied brain regions are the amygdala (Amyg, 17%), hippocampus (HPF, 20%) and medial prefrontal cortex (mPFC, 38%). HPF and mPFC have been implicated in both negative and positive affect and have each been tested in fear-like, anxiety-like and depressive-like behaviors. Amyg has mainly been tested in the context of fear where, consistent with early lesion studies, it is observed to primarily mediate negative affect. Despite a variety of efferent and afferent Amyg pathways being evaluated in anxiety- and depressive-like behaviors, no region-specific perturbation has evaluated its role in these behaviors. Of the 47 total region focused studies, 25% targeted molecularly identified subpopulations of neurons, with the majority targeting excitatory neurons with a Calcium calmodulin-dependent protein kinase II (CamKII) promoter.

A total of 60 studies targeting 65 unique pathways (including bidirectional control) were identified (3 pathway studies overlapped with targeting of particular brain regions, **Table 4**). Seventeen studies targeted more two pathways and three studies targeted three or more pathways. The majority (58%) targeted a specific cell type: 46% utilized CamKII to target excitatory neurons, 6% targeted somatostatin (SST)-expressing neurons, 3.4% targeted tyrosine hydroxylase (TH)-positive neurons, and 3.3% targeted parvalbumin (PV)-positive neurons. Dopamine (D2)-, vesicular glutamate (Vglut)-, serotonin (5HT)-, and corticotropin releasing factor (CRF)-expressing neurons were each targeted once (1.6% each). Of the 65 pathways, only eleven have been manipulated in more than one study. Ten of these pathways were investigated by different labs with three pathways showing incongruent results. Some inconsistencies are due to targeting different neuronal cell-types. Studies demonstrating increased negative affective-like behaviors with HPF to mPFC pathway (HPF→mPFC) stimulation, targeted excitatory neurons (27–29). On the other hand, studies demonstrating decreased negative affective-like behaviors with HPF→mPFC stimulation used a pan-neuronal promoter (30,31). Others are due to behavioral differences: HPF→Amyg is implicated in opposing affective valence, but negative affect is promoted in fear-related behavior (32,33) and positive affect is promoted in depression-related behaviors (34). Lastly, the ventral tegmental area to nucleus accumbens pathway (VTA→ACB) increases depressive-like behaviors with phasic stimulation (35,36), but decreases these behaviors with other stimulation patterns (37). Very few pathways display consistencies in both directions. For example, both the infralimbic and prelimibic connection (IL←→PL) (38,39) and the PL←→BLA (9, 62, 64) connection have been demonstrated to reliably increase fear-like behaviors. Implication of Amyg→HPF in increased depressive- and anxiety-like behaviors has been replicated across multiple studies, but the pathway was targeted exclusively by one research team, likely contributing to the reproducibility (40,41). In summary, most pathways have been investigated in single studies by unique teams with very few replicated across multiple studies and/or laboratories.

**Table 4.**
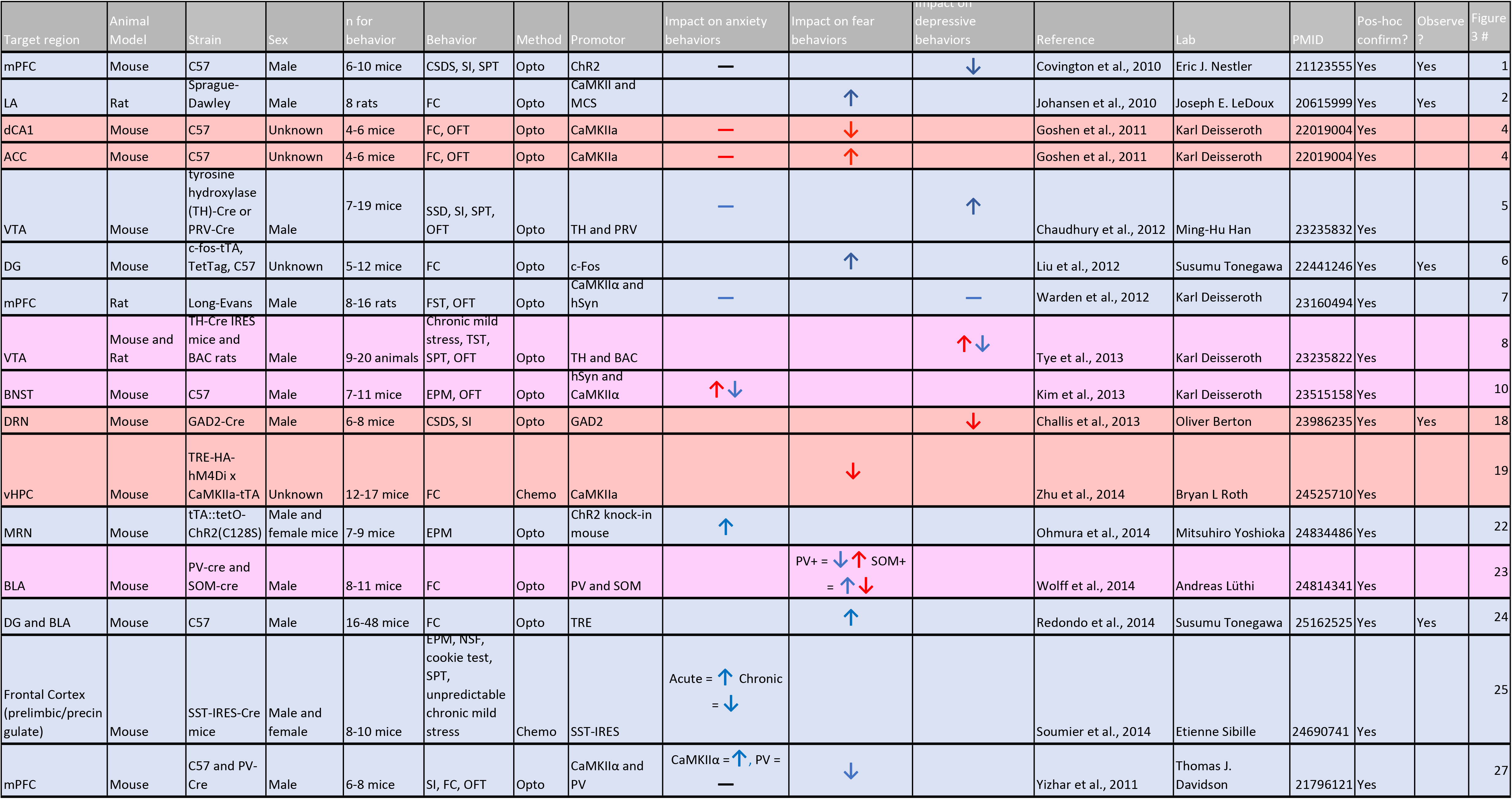
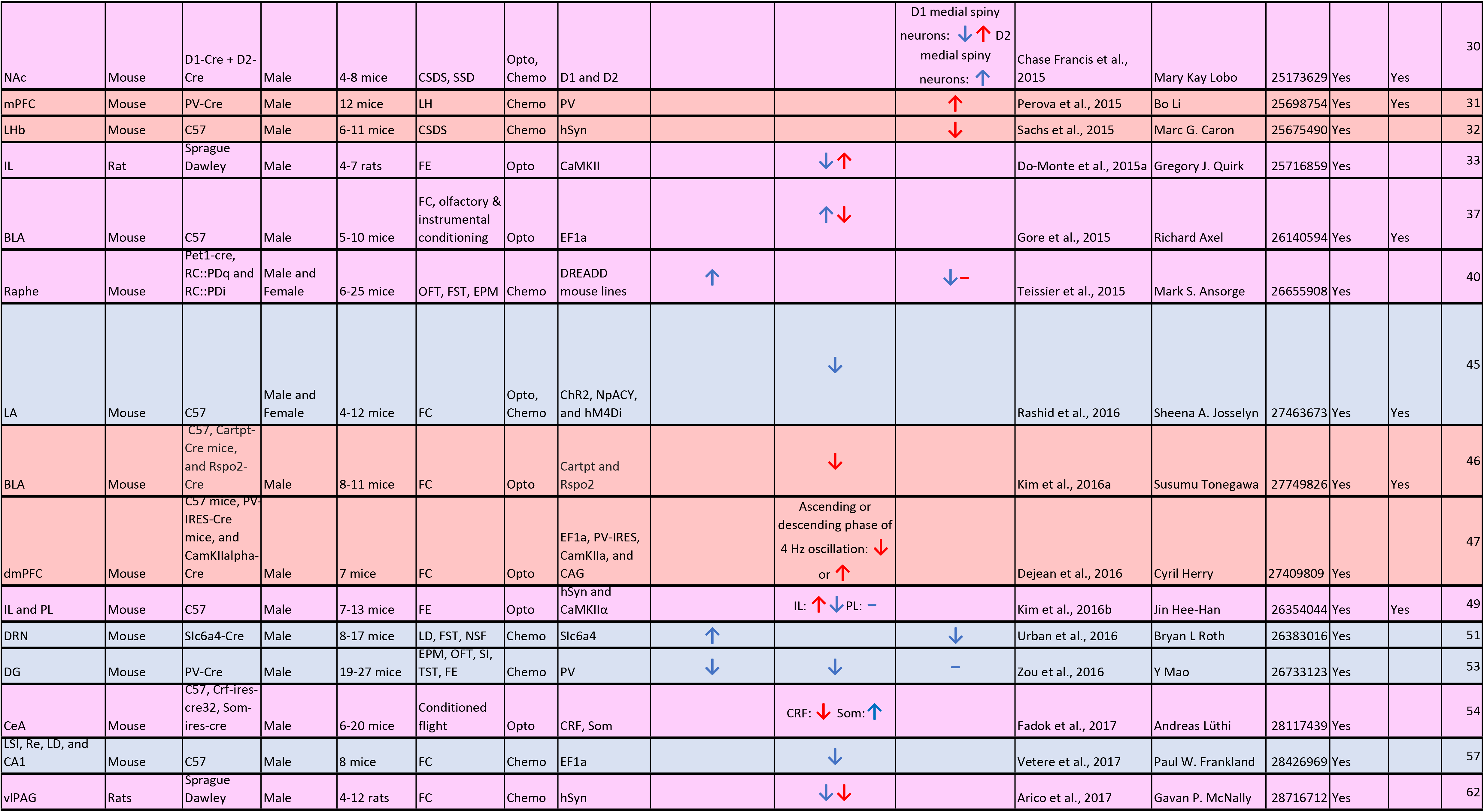
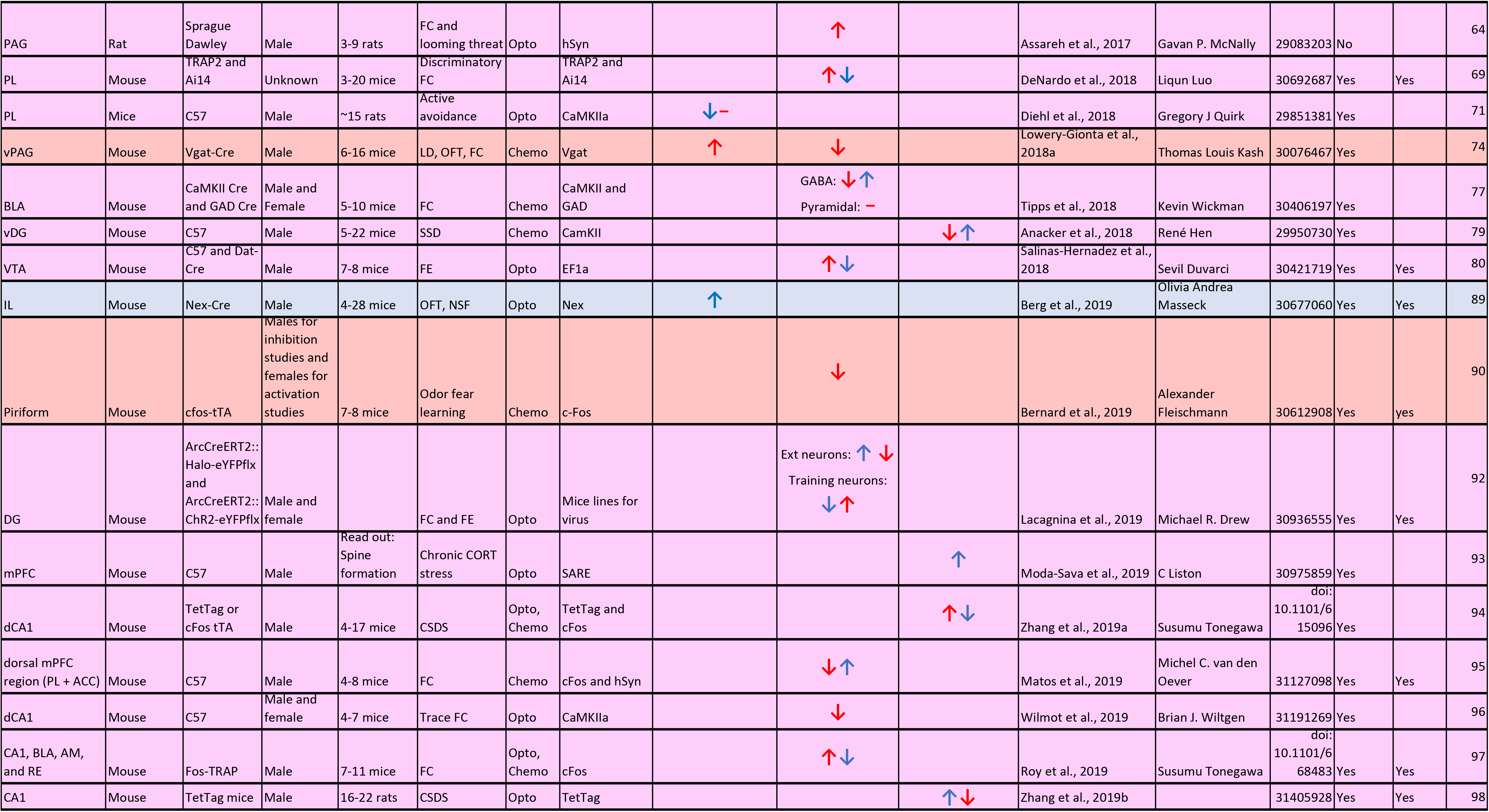
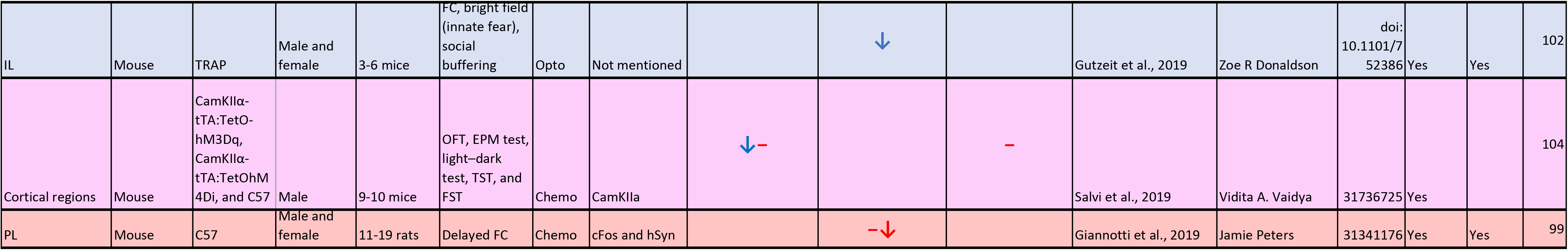
Region Focused. Details from studies focused on dissecting the role of a region in a perturbed affective-like state. Blue rows/arrows represent activation, red rows/arrows represent inhibition, purple rows indicate both activation and inhibition.

| Target region | Animal Model | Strain | Sex | n for behavior | Behavior | Method | Promotor | Impact on anxiety behaviors | Impact on fear behaviors | Impact on depressive behaviors | Reference | Lab | PMID | Pos-hoc confirm? | Observed? | Figure 3 # |
| --- | --- | --- | --- | --- | --- | --- | --- | --- | --- | --- | --- | --- | --- | --- | --- | --- |
| mPFC | Mouse | C57 | Male | 6-10 mice | CSDS, SI, SPT | Opto | Chr2 | — |  | ↓ | Covington et al., 2010 | Eric J. Nestler | 21123555 | Yes | Yes | 1 |
| LA | Rat | Sprague-Dawley | Male | 8 rats | FC | Opto | CaMKII and MCS |  | ↑ |  | Johansen et al., 2010 | Joseph E. LeDoux | 20615999 | Yes | Yes | 2 |
| dCA1 | Mouse | C57 | Unknown | 4-6 mice | FC, OFT | Opto | CaMKIIa | — | ↓ |  | Goshen et al., 2011 | Karl Deisseroth | 22019004 | Yes |  | 4 |
| ACC | Mouse | C57 | Unknown | 4-6 mice | FC, OFT | Opto | CaMKIIa | — | ↑ |  | Goshen et al., 2011 | Karl Deisseroth | 22019004 | Yes |  | 4 |
| VTA | Mouse | tyrosine hydroxylase (TH)-Cre or PRV-Cre | Male | 7-19 mice | SSD, SI, SPT, OFT | Opto | TH and PRV | — |  | ↑ | Chaudhury et al., 2012 | Ming-Hu Han | 23235832 | Yes |  | 5 |
| DG | Mouse | c-fos-tTA, TetTag, C57 | Unknown | 5-12 mice | FC | Opto | c-Fos |  | ↑ |  | Liu et al., 2012 | Susumu Tonegawa | 22441246 | Yes | Yes | 6 |
| mPFC | Rat | Long-Evans | Male | 8-16 rats | FST, OFT | Opto | CaMKIIα and hSyn | — |  | — | Warden et al., 2012 | Karl Deisseroth | 23160494 | Yes |  | 7 |
| VTA | Mouse and Rat | TH-Cre IRES mice and BAC rats | Male | 9-20 animals | Chronic mild stress, TST, SPT, OFT | Opto | TH and BAC |  |  | ↑↓ | Tye et al., 2013 | Karl Deisseroth | 23235822 | Yes |  | 8 |
| BNST | Mouse | C57 | Male | 7-11 mice | EPM, OFT | Opto | hSyn and CaMKIIα | ↑↓ |  |  | Kim et al., 2013 | Karl Deisseroth | 23515158 | Yes |  | 10 |
| DRN | Mouse | GAD2-Cre | Male | 6-8 mice | CSDS, SI | Opto | GAD2 |  |  | ↓ | Challis et al., 2013 | Oliver Berton | 23986235 | Yes | Yes | 18 |
| vHPC | Mouse | TRE-HA-hM4Di x CaMKIIa-tTA | Unknown | 12-17 mice | FC | Chemo | CaMKIIa |  | ↓ |  | Zhu et al., 2014 | Bryan L Roth | 24525710 | Yes |  | 19 |
| MRN | Mouse | tTA::tetO-ChR2(C128S) | Male and female mice | 7-9 mice | EPM | Opto | Chr2 knock-in mouse | ↑ |  |  | Ohmura et al., 2014 | Mitsuhiro Yoshioka | 24834486 | Yes |  | 22 |
| BLA | Mouse | PV-cre and SOM-cre | Male | 8-11 mice | FC | Opto | PV and SOM |  | PV+ = ↓↑ SOM+ = ↑↓ |  | Wolff et al., 2014 | Andreas Lüthi | 24814341 | Yes |  | 23 |
| DG and BLA | Mouse | C57 | Male | 16-48 mice | FC | Opto | TRE |  | ↑ |  | Redondo et al., 2014 | Susumu Tonegawa | 25162525 | Yes | Yes | 24 |
| Frontal Cortex (prelimbic/precingulate) | Mouse | SST-IRES-Cre mice | Male and female | 8-10 mice | EPM, NSF, cookie test, SPT, unpredictable chronic mild stress | Chemo | SST-IRES | Acute = ↑ Chronic = ↓ |  |  | Soumier et al., 2014 | Etienne Sibille | 24690741 | Yes |  | 25 |
| mPFC | Mouse | C57 and PV-Cre | Male | 6-8 mice | SI, FC, OFT | Opto | CaMKIIα and PV | CaMKIIα = ↑, PV = — | ↓ |  | Yizhar et al., 2011 | Thomas J. Davidson | 21796121 | Yes |  | 27 |
| NAc | Mouse | D1-Cre + D2-Cre | Male | 4-8 mice | CSDS, SSD | Opto, Chemo | D1 and D2 |  |  | D1 medial spiny neurons: ↓↑ D2 medial spiny neurons: ↑ | Chase Francis et al., 2015 | Mary Kay Lobo | 25173629 | Yes | Yes | 30 |
| mPFC | Mouse | PV-Cre | Male | 12 mice | LH | Chemo | PV |  |  | ↑ | Perova et al., 2015 | Bo Li | 25698754 | Yes | Yes | 31 |
| LHb | Mouse | C57 | Male | 6-11 mice | CSDS | Chemo | hSyn |  |  | ↓ | Sachs et al., 2015 | Marc G. Caron | 25675490 | Yes |  | 32 |
| IL | Rat | Sprague Dawley | Male | 4-7 rats | FE | Opto | CaMKII |  | ↓↑ |  | Do-Monte et al., 2015a | Gregory J. Quirk | 25716859 | Yes |  | 33 |
| BLA | Mouse | C57 | Male | 5-10 mice | FC, olfactory & instrumental conditioning | Opto | EF1a |  | ↑↓ |  | Gore et al., 2015 | Richard Axel | 26140594 | Yes | Yes | 37 |
| Raphe | Mouse | Pet1-cre, RC::PDq and RC::PDi | Male and Female | 6-25 mice | OFT, FST, EPM | Chemo | DREADD mouse lines | ↑ |  | ↓- | Teissier et al., 2015 | Mark S. Ansorge | 26655908 | Yes |  | 40 |
| LA | Mouse | C57 | Male and Female | 4-12 mice | FC | Opto, Chemo | Chr2, NpACY, and hM4Di |  | ↓ |  | Rashid et al., 2016 | Sheena A. Josselyn | 27463673 | Yes | Yes | 45 |
| BLA | Mouse | C57, Cartpt-Cre mice, and Rspo2-Cre | Male | 8-11 mice | FC | Opto | Cartpt and Rspo2 |  | ↓ |  | Kim et al., 2016a | Susumu Tonegawa | 27749826 | Yes | Yes | 46 |
| dmPFC | Mouse | C57 mice, PV-IRES-Cre mice, and CamKIIalpha-Cre | Male | 7 mice | FC | Opto | EF1a, PV-IRES, CamKIIa, and CAG |  | Ascending or descending phase of 4 Hz oscillation: ↓ or ↑ |  | Dejean et al., 2016 | Cyril Herry | 27409809 | Yes |  | 47 |
| IL and PL | Mouse | C57 | Male | 7-13 mice | FE | Opto | hSyn and CaMKIIα |  | IL: ↑↓ PL: - |  | Kim et al., 2016b | Jin Hee-Han | 26354044 | Yes | Yes | 49 |
| DRN | Mouse | Slc6a4-Cre | Male | 8-17 mice | LD, FST, NSF | Chemo | Slc6a4 | ↑ |  | ↓ | Urban et al., 2016 | Bryan L Roth | 26383016 | Yes |  | 51 |
| DG | Mouse | PV-Cre | Male | 19-27 mice | EPM, OFT, SI, TST, FE | Chemo | PV | ↓ | ↓ | - | Zou et al., 2016 | Y Mao | 26733123 | Yes |  | 53 |
| CeA | Mouse | C57, Crf-ires-cre32, Som-ires-cre | Male | 6-20 mice | Conditioned flight | Opto | CRF, Som |  | CRF: ↓ Som: ↑ |  | Fadok et al., 2017 | Andreas Lüthi | 28117439 | Yes |  | 54 |
| LSI, Re, LD, and CA1 | Mouse | C57 | Male | 8 mice | FC | Chemo | EF1a |  | ↓ |  | Vetere et al., 2017 | Paul W. Frankland | 28426969 | Yes |  | 57 |
| vIPAG | Rats | Sprague Dawley | Male | 4-12 rats | FC | Chemo | hSyn |  | ↓↓ |  | Arico et al., 2017 | Gavan P. McNally | 28716712 | Yes |  | 62 |
| PAG | Rat | Sprague Dawley | Male | 3-9 rats | FC and looming threat | Opto | hSyn |  | ↑ |  | Assareh et al., 2017 | Gavan P. McNally | 29083203 | No |  | 64 |
| PL | Mouse | TRAP2 and Ai14 | Unknown | 3-20 mice | Discriminatory FC |  | TRAP2 and Ai14 |  | ↑↓ |  | DeNardo et al., 2018 | Liqun Luo | 30692687 | Yes | Yes | 69 |
| PL | Mice | C57 | Male | ~15 rats | Active avoidance | Opto | CaMKIIa | ↓- |  |  | Diehl et al., 2018 | Gregory J Quirk | 29851381 | Yes |  | 71 |
| vPAG | Mouse | Vgat-Cre | Male | 6-16 mice | LD, OFT, FC | Chemo | Vgat | ↑ | ↓ |  | Lowery-Gionta et al., 2018a | Thomas Louis Kash | 30076467 | Yes |  | 74 |
| BLA | Mouse | CaMKII Cre and GAD Cre | Male and Female | 5-10 mice | FC | Chemo | CaMKII and GAD |  | GABA: ↓↑<br>Pyramidal: - |  | Tipps et al., 2018 | Kevin Wickman | 30406197 | Yes |  | 77 |
| vDG | Mouse | C57 | Male | 5-22 mice | SSD | Chemo | CamKII |  |  | ↓↑ | Anacker et al., 2018 | René Hen | 29950730 | Yes |  | 79 |
| VTA | Mouse | C57 and Dat-Cre | Male | 7-8 mice | FE | Opto | EF1a |  | ↑↓ |  | Salinas-Hernandez et al., 2018 | Sevil Duvarci | 30421719 | Yes | Yes | 80 |
| IL | Mouse | Nex-Cre | Male | 4-28 mice | OFT, NSF | Opto | Nex | ↑ |  |  | Berg et al., 2019 | Olivia Andrea Masseck | 30677060 | Yes | Yes | 89 |
| Piriform | Mouse | cfos-tTA | Males for inhibition studies and females for activation studies | 7-8 mice | Odor fear learning | Chemo | c-Fos |  | ↓ |  | Bernard et al., 2019 | Alexander Fleischmann | 30612908 | Yes | yes | 90 |
| DG | Mouse | ArcCreERT2::Halo-eYFPflx and ArcCreERT2::Chr2-eYFPflx | Male and female |  | FC and FE | Opto | Mice lines for virus |  | Ext neurons: ↑↓<br>Training neurons: ↓↑ |  | Lacagnina et al., 2019 | Michael R. Drew | 30936555 | Yes | Yes | 92 |
| mPFC | Mouse | C57 | Male | Read out: Spine formation | Chronic CORT stress | Opto | SARE |  |  | ↑ | Moda-Sava et al., 2019 | C Liston | 30975859 | Yes |  | 93 |
| dCA1 | Mouse | TetTag or cFos tTA | Male | 4-17 mice | CSDS | Opto, Chemo | TetTag and cFos |  |  | ↑↓ | Zhang et al., 2019a | Susumu Tonegawa | doi: 10.1101/615096 | Yes |  | 94 |
| dorsal mPFC region (PL + ACC) | Mouse | C57 | Male | 4-8 mice | FC | Chemo | cFos and hSyn |  | ↓↑ |  | Matos et al., 2019 | Michel C. van den Oever | 31127098 | Yes | Yes | 95 |
| dCA1 | Mouse | C57 | Male and female | 4-7 mice | Trace FC | Opto | CaMKIIa |  | ↓ |  | Wilmot et al., 2019 | Brian J. Wiltgen | 31191269 | Yes |  | 96 |
| CA1, BLA, AM, and RE | Mouse | Fos-TRAP | Male | 7-11 mice | FC | Opto, Chemo | cFos |  | ↑↓ |  | Roy et al., 2019 | Susumu Tonegawa | doi: 10.1101/668483 | Yes | Yes | 97 |
| CA1 | Mouse | TetTag mice | Male | 16-22 rats | CSDS | Opto | TetTag |  |  | ↑↓ | Zhang et al., 2019b |  | 31405928 | Yes | Yes | 98 |
| IL | Mouse | TRAP | Male and female | 3-6 mice | FC, bright field (innate fear), social buffering | Opto | Not mentioned |  | ↓ |  | Gutzeit et al., 2019 | Zoe R Donaldson | doi: 10.1101/752386 | Yes | Yes | 102 |
| Cortical regions | Mouse | CamKIIα-tTA:TetO-hM3Dq, CamKIIα-tTA:TetOhM4Di, and C57 | Male | 9-10 mice | OFT, EPM test, light-dark test, TST, and FST | Chemo | CamKIIa | ↓ - |  | - | Salvi et al., 2019 | Vidita A. Vaidya | 31736725 | Yes |  | 104 |
| PL | Mouse | C57 | Male and female | 11-19 rats | Delayed FC | Chemo | cFos and hSyn |  | - ↓ |  | Giannotti et al., 2019 | Jamie Peters | 31341176 | Yes | Yes | 99 |

**Table 5.**
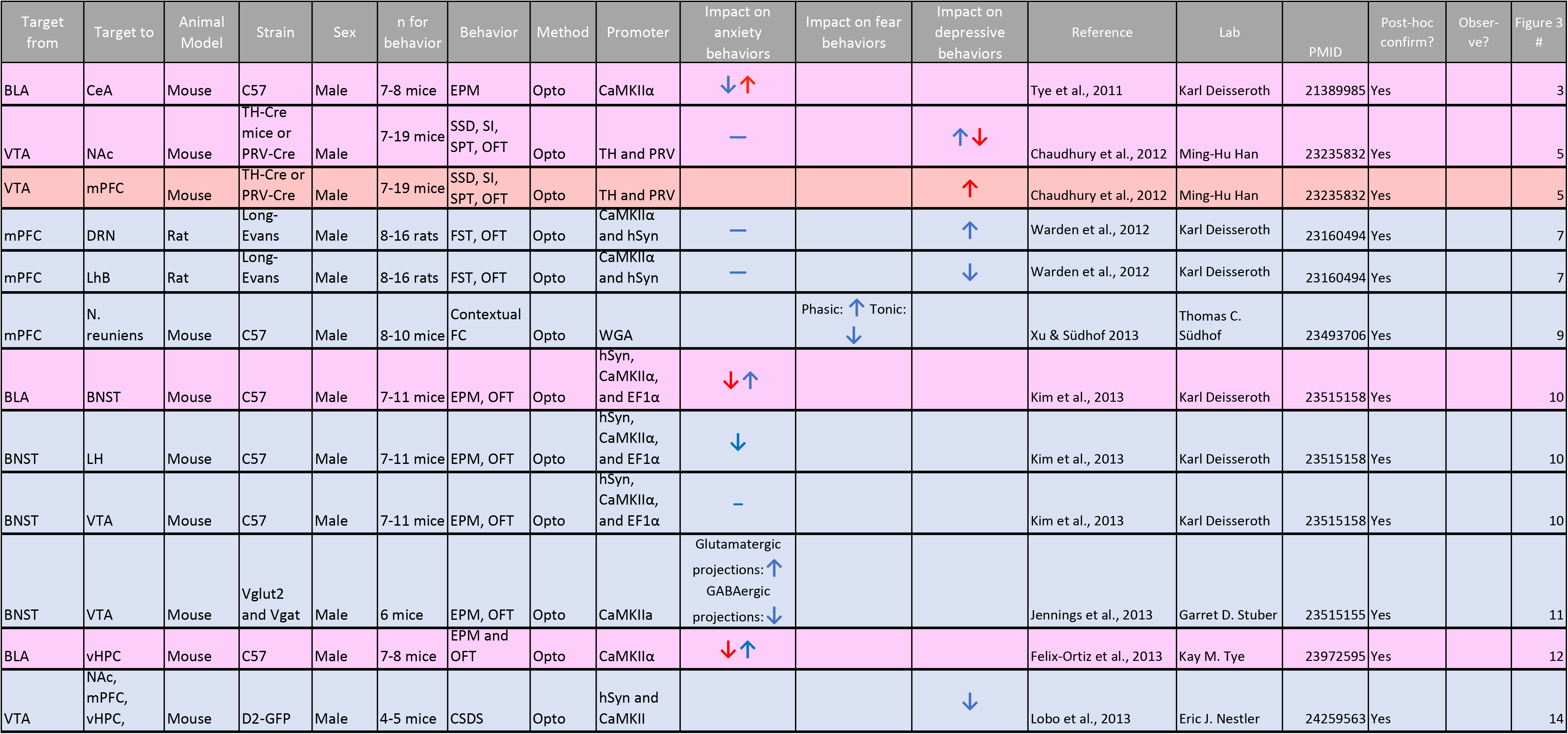
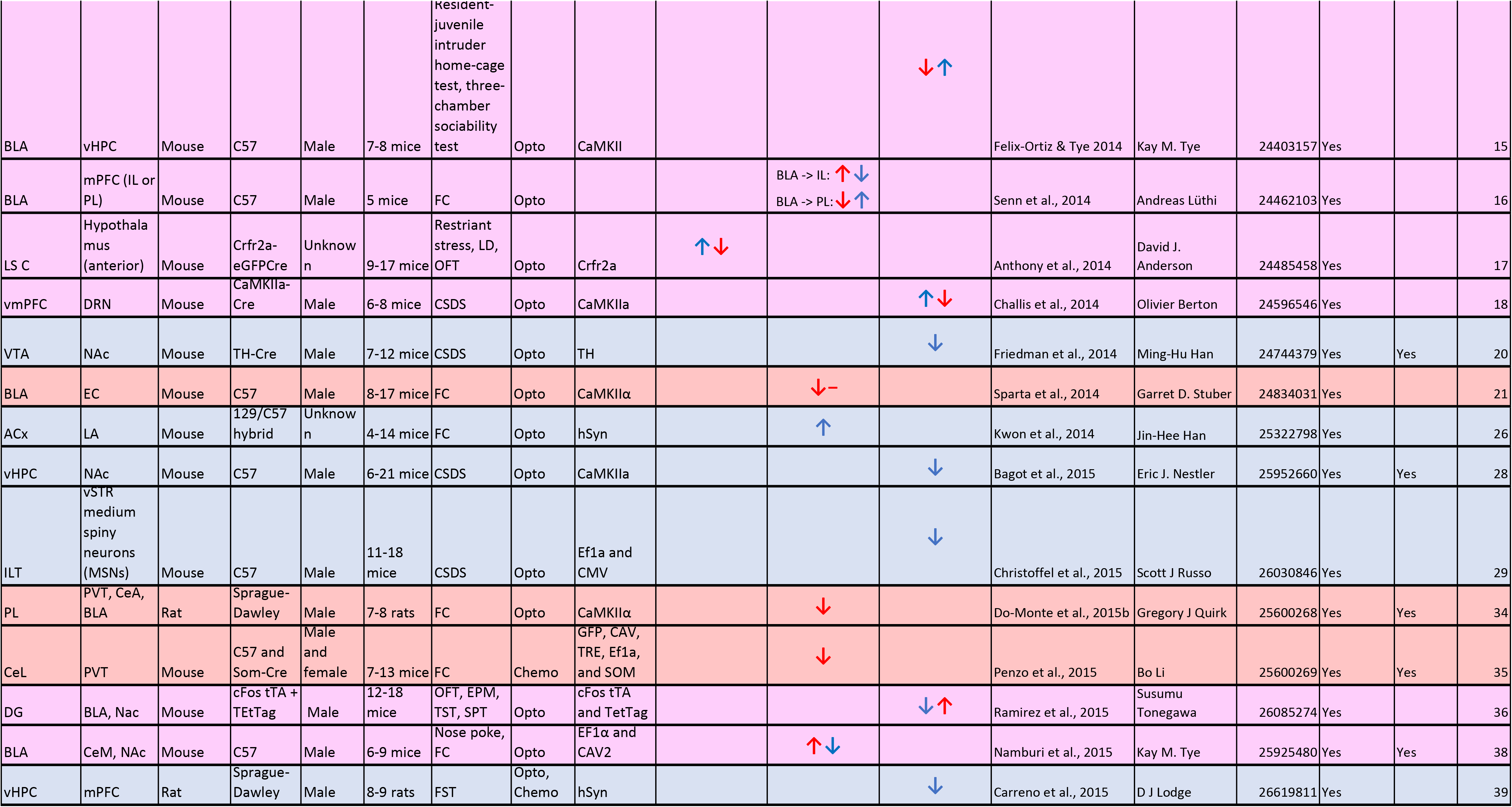
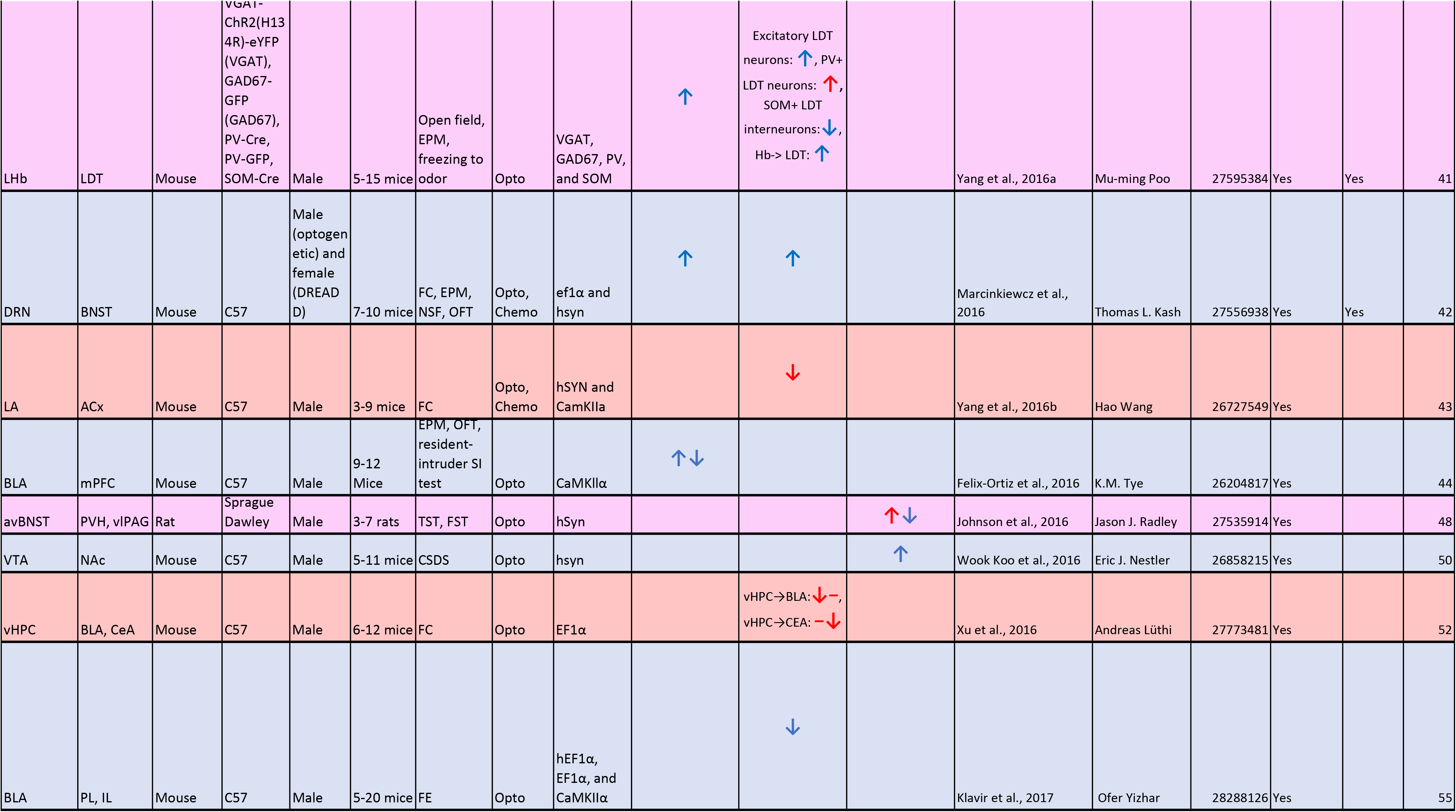
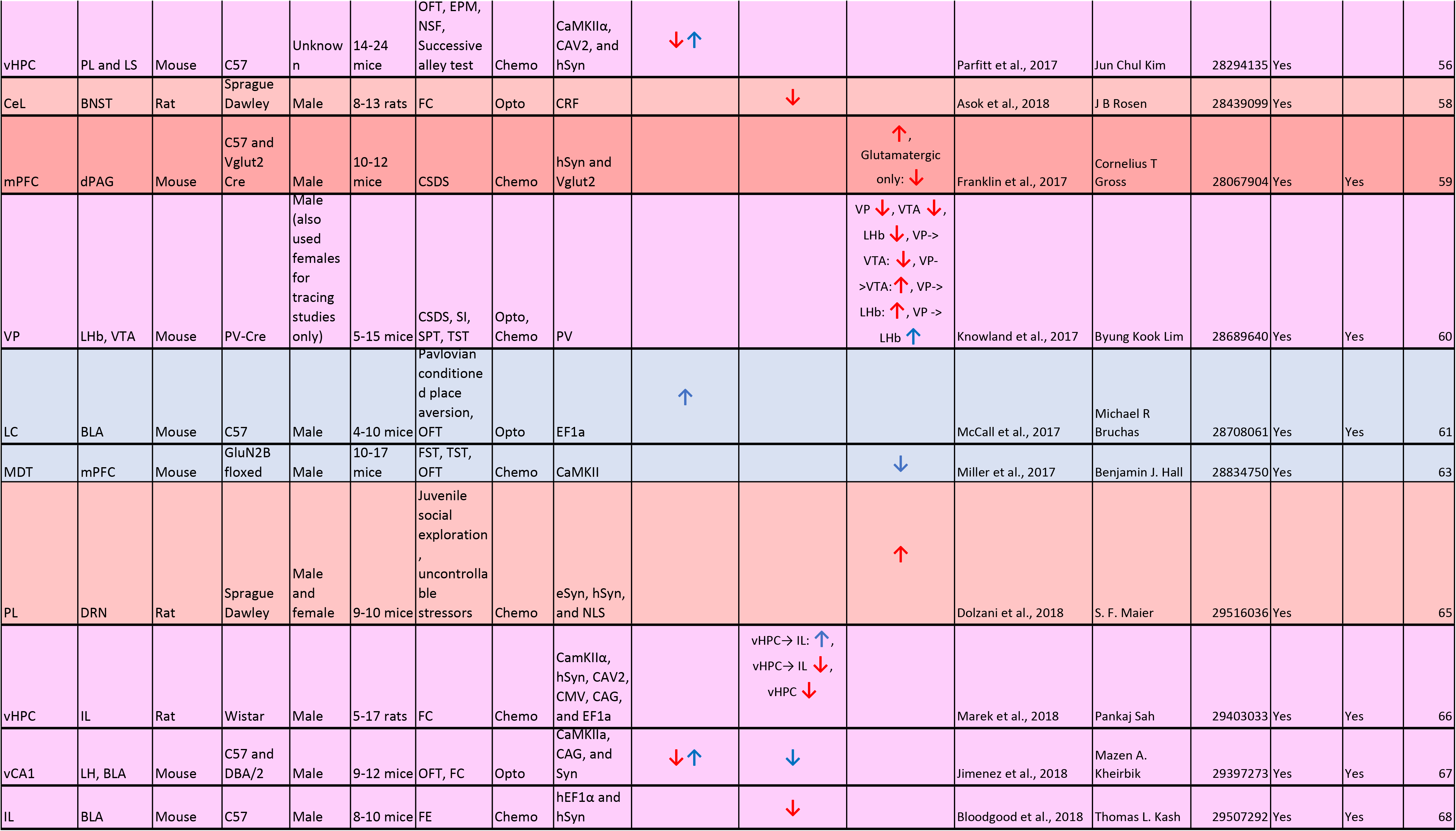
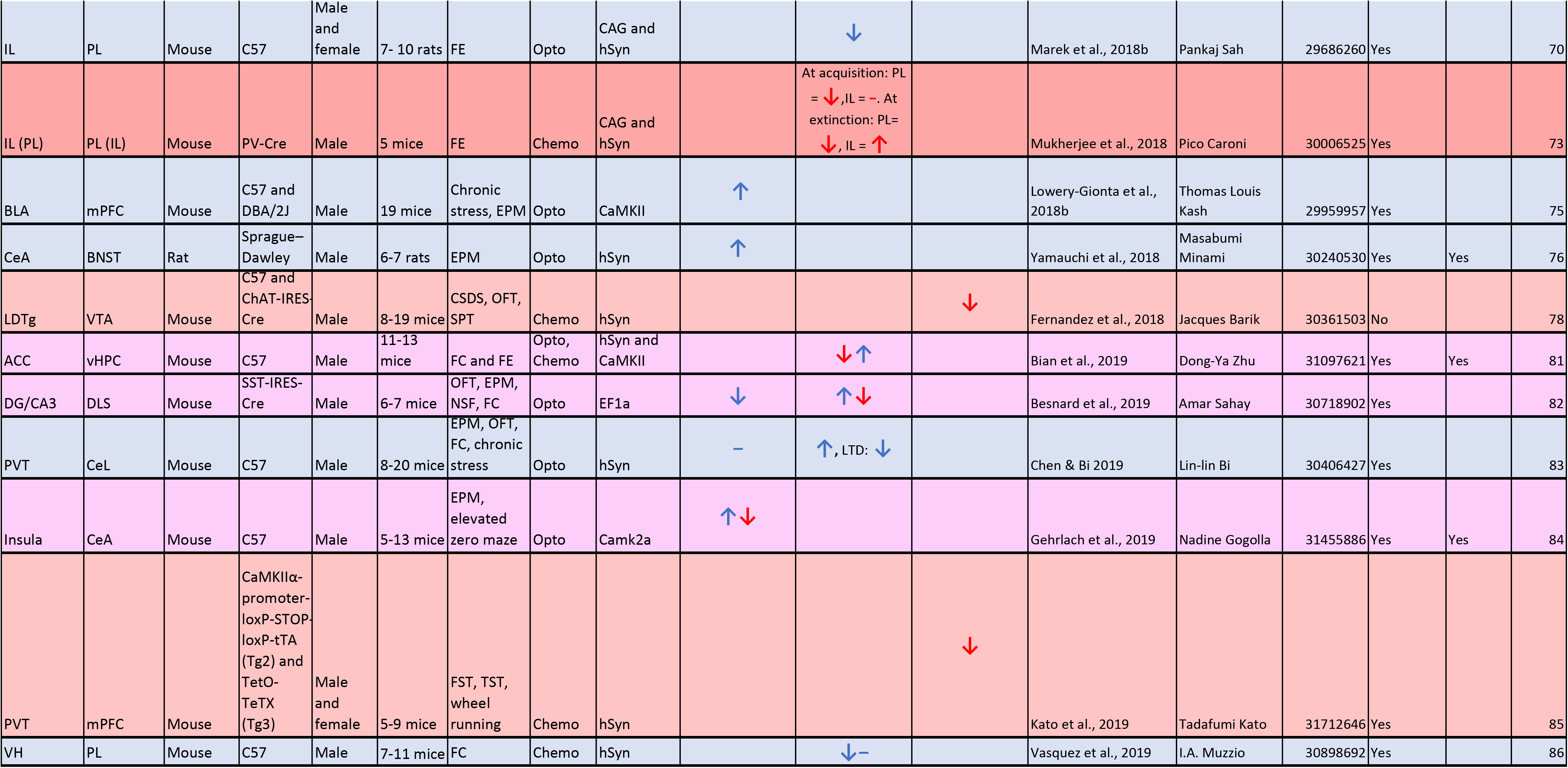
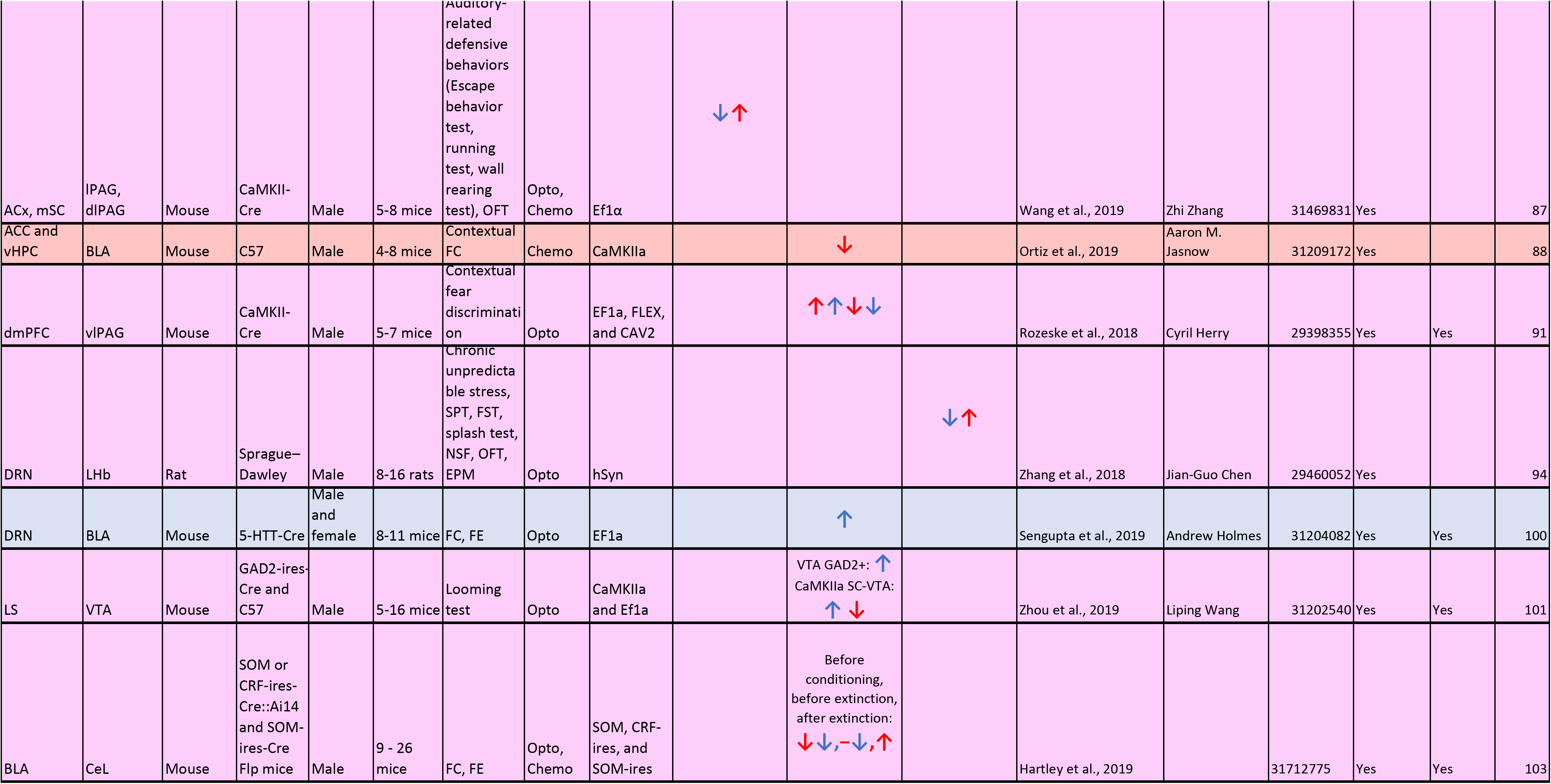
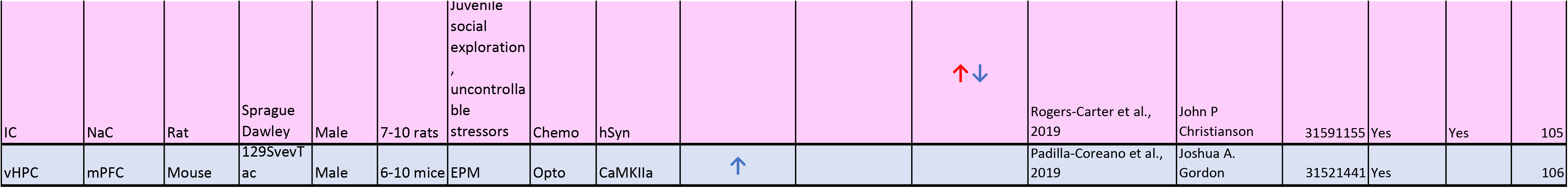
Pathway-focused. Details from studies focused on dissecting the role of a pathway in a perturbed affective-like state Blue rows/arrows represent activation, red rows/arrows represent inhibition, purple rows indicate both activation and inhibition.

More than half the accumulated data (overall 62%, 34/60 pathway focused and 32/49 region focused) has been obtained with circuit perturbations during a single behavioral test. Very few studies have investigated a pathway across multiple affective-like domains (overall 12%, 7/59 pathway focused, 6/47 region focused). This generally entailed testing anxiety-like behavior using open field and fear-like behavior using fear conditioning, and mostly implicated the manipulated circuit in one and not the other behavior (42–44). Pathways investigated multiple times have primarily only been dissected with one tool; for example, both VTA→ACB (35–37) and Amyg→bed nucleus of the stria terminalis (BST) (45–47) have only been studied with optogenetics.

Surprisingly, a minority of studies (40%) conducted some type of observation of the targeted circuit in an endogenous state prior to perturbation (21/60 pathway focused and 22/49 region focused). A loose definition of observation was applied, including checking that an IEG is expressed in the region or pathway of interest after behavior, using tracing to ensure a functional connection exists between two regions, and utilizing a “trapping” technology to indelibly mark neurons activated by a particular behavior (e.g. Tet/Tag (48) or TRAP (49) mice). After removal of studies using trapping techniques, only 22% report an observation of endogenous dynamics of a circuit prior to circuit manipulation.

## CONNECTOMIC PRINCIPLES OF RODENT AFFECTIVE CIRCUITRY

### Historical perspective

When visualized over time, a historical perspective emerges (**Supplementary video 1**). Initial studies targeted unitary regions. Studies tracing pathways to and from these identified nodes followed. Most recently, there was a return to regional analysis, motivated by cell-type specific circuit dissection. Initial studies focused heavily on the triad of brain regions strongly implicated in human mood disorders: Amyg, HPF, and mPFC (50–52). Over time, analysis expanded to regions further and further removed from this triad, some of which have less obvious ties to human conditions. For example, the Nucleus reuniens (NR) has not directly been investigated in human affective states. A PubMed search (accessed on December 30th, 2019) for ‘N. reuniens’, ‘human’, and ‘affect’ (or ‘anxiety’, ‘fear’, and ‘depression’) collectively only reveals one paper, which reviews the overall role of the thalamus and all its sub-regions in animal behaviors, both affective and cognitive (53). Therefore, although there are lines of evidence for the role of this nucleus in fear-like behavioral regulation in animal models (54), there is little evidence of its relevance to human mental health disorders.

Viewed as a whole, the cumulative connectome also conjures some obvious holes. The cerebellum for example, has recently become implicated in mood disorders in both humans (55–58) and rodents (59). The cerebellum sends and receives inputs from numerous cortical regions (60,61), yet neither regional nor pathway analysis has to-date probed cerebellar contribution to rodent affective-like behaviors.

### Node centrality

The centrality of the Amyg, and its primary role of mediating negative affect (red lines in **Figure 3**), is immediately apparent. The mPFC is another notable node, mediating both positive and negative affect depending on the specific pathway or subregion targeted. The bidirectional interaction between the mPFC and Amyg as a mediator of negative affect is well-replicated among multiple labs using multiple paradigms (illustrated by the thicker red line in **Figure 3**, which is proportional to the number of studies evaluating this pathway) (9,62–65).

When fear-, anxiety-, or depressive-like circuits are visualized separately **(Figure 4)**, unique patterns emerge. The mPFC dominates the network for depressive-like behaviors, the amygdala dominates the fear circuit, and anxiety-like behaviors are mediated by a more distributed network.

**Fig. 4.**
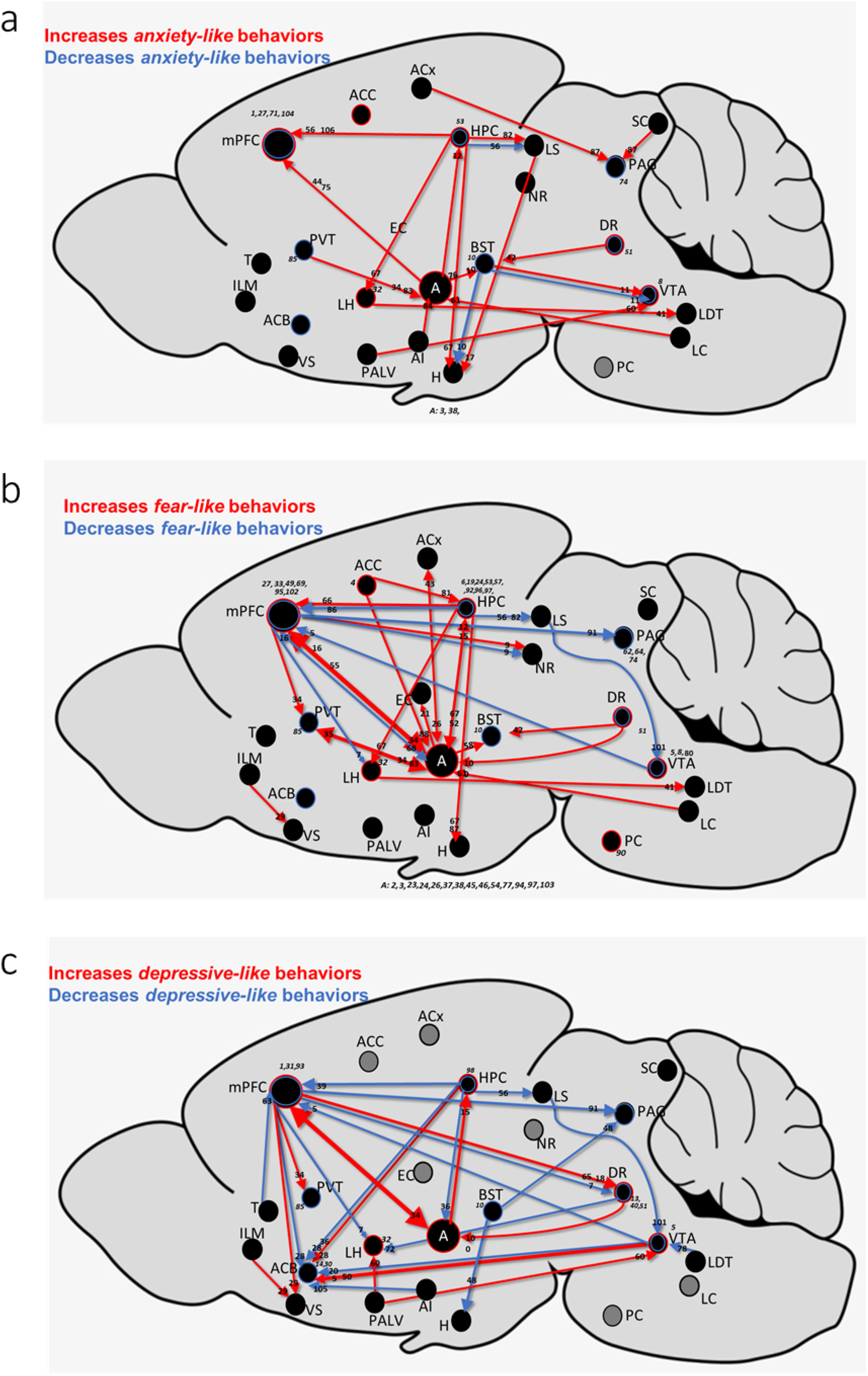
Verified rodent brain functional connectivity in specific affective states: anxiety-, fear, and depressive-like. Pathway and region manipulations sorted by the behavioral paradigm that was paired with the optogenetic or chemogenetic manipulation. (a) Anxiety-like behaviors presented in the backdrop of the whole rodent brain. (b) Fear-like behaviors presented in the backdrop of the whole rodent brain. (c) Depressive-like behaviors presented in the backdrop of the whole rodent brain. Red indicates regions and pathways in which activation promotes negative affective-like behavior, while blue indicates regions and pathways in which activation promotes positive

It is important to consider that this node centrality emerged in a hypothesis-driven context. Thus, despite the cumulative evidence identifying Amyg as a “fear center” (and the preceding decades of non-circuit-based studies investigating this region’s role in fear), none of the studies reviewed performed any type of brain-wide approach to first confirm the region’s central role to the particular behavior studied. While this is intuitive given the stepwise progress expected of scientific inquiry, it is nevertheless an important observation that needs careful scrutiny. The danger of circuit era mapping is that, as each individual pathway is added to the rodent affective-like connectome, the resulting network structure could move further and further away from the “ground-truth” connectome associated with a particular behavior due to propagation of error (66).

### Illuminating regional and subregional specificity

The multi-leveled specificity of circuit era tools (**Figure 1**) has led to increasingly refined understanding of regional and subregional contributions to behavioral outcomes. The first layer of specificity is imparted by viral injections, which offer improved anatomical localization over lesion and pharmacological studies (9,67). A secondary layer of specificity can then be added using cell-type specific gene-expression. As an example, serotonergic neurons are anatomically restricted to the raphe nuclei, tiny regions in the mid-hindbrain traditionally difficult to precisely target. To achieve high-degree of regional specificity, studies of raphe nuclei (dorsal raphe nucleus [DR] and/or medial raphe nucleus [MRN]) commonly restrict expression of opsins or DREADDs to serotonergic neurons using the tryptophan hydroxylase 2 (Tph2) promoter (68), the fifth Ewing variant (FEV) promoter (69), or the SIc6a4 gene (128). It should be noted however, that while such genetic labeling techniques impart high-degree of anatomical specificity, they can also potentially miss key aspects of functioning. Serotonergic neurons actually only make up 20% of all neurons in the median raphe region, with glutamatergic and GABAergic neurons predominating (71). Elucidation of a region’s function should ideally include both regional specificity as well as an understanding of the interplay between various neuronal types. Of the four re viewed studies of DR/MRN, one began addressing this issue by specifically targeting GABAergic neurons in the region (70).

Finally, intersectional approaches impart projection-specificity. This is achieved by injecting a retrograde virus carrying a recombinase, such as Cre, in an efferent region in combination with a second injection of a virus carying Cre-dependent opsins or DREADDs into the region of interest. This results in selective targeting of neurons based on their axonal projections.

**Figure 5** shows the detailed subregional data generated using these approaches on two key brain areas involved in affective-like behaviors: the mPFC and the basolateral amygdala (BLA). The IL, a tiny subregion of the mPFC, is a great example of the technological advances ushered in with circuit era tools. Lesion and electrophysiological studies had previously provided contradicting data on the role of IL in fear extinction, with some studies implicating IL in fear extinction (72) and others reporting IL lesions not to impair extinction and IL firing not to be associated with extinction (72,73). Circuit manipulations, thus far, have unequivocally demonstrated that functional activation of the IL plays a crucial role in extinction learning (9,38,53,54,55,62,64,67,75). In contrast, the neighboring PL subregion of the mPFC has been implicated in fear memory formation (74–76).

**Fig. 5.**
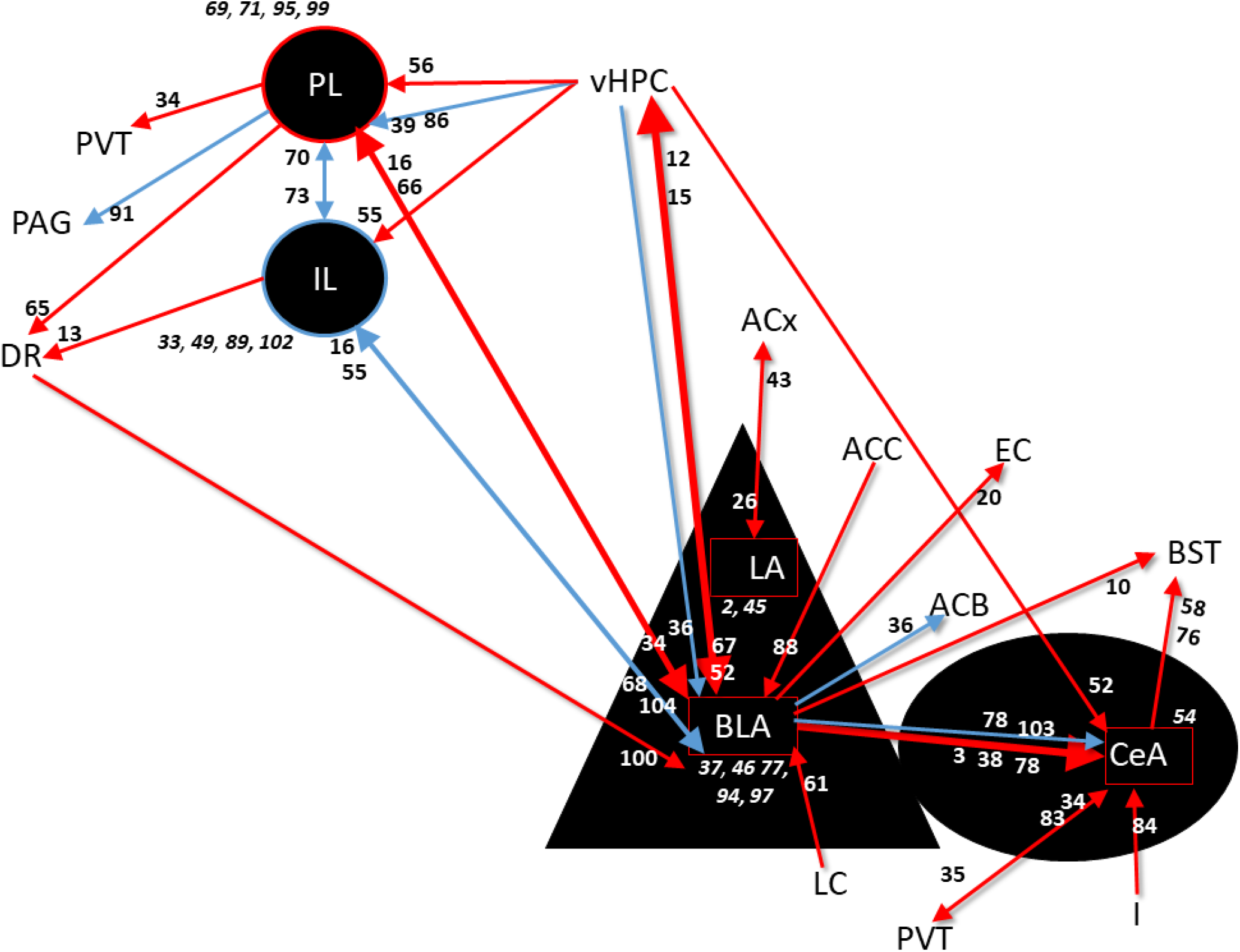
Detailed dissection of subregional contributions to affective-like behaviors in two commonly studied regions. Zoom in on the medial prefrontal cortex, showing the opposing contributions of the prelimbic (PL) and infralimbic (IL) subregions to affective-like behaviors. Zoom in on the amygdala and its subregions: lateral amygdala (LA), basolateral amygdala (BLA), central amygdala(CeA). All studies represent optogenetic or chemogenetic studies coupled with rodent models of disordered affect.

Circuit era tools have also refined our understanding of different circuits within the BLA. This region contains various types of neurons that play distinctive roles in fear processing. Initially, this was appreciated using electrophysiological approaches that identified “fear neurons”, responding to fear learning, and “fear extinction” neurons, responding to extinction learning (77). Interestingly, these neurons also display different connectivity patterns, with fear neurons preferentially receiving inputs from the ventral hippocampus (vHPC) and extinction neurons from the mPFC (77), stressing the necessity of both structural and functional specificity in circuit manipulations. Following up on these experiments, optogenetic stimulation confirmed the vHPC →BLA pathway’s involvement in fear memory formation (32,78), and chemogenetic inhibition confirmed the importance of the IL→BLA pathway in extinction learning (95). Distinct roles in fear learning have also been observed in various neuronal subtypes in the BLA. Activation of PV interneurons during a conditioned stimulus presentation promotes auditory fear learning whereas activation of SST interneurons inhibits learning (80).

### The final frontier: activity-dependent targeting of circuits

The final frontier in specificity has taken advantage of immediate early genes (IEGs) to target neurons activated during particular behaviors, thereby adding temporal specificity. Memories and behavior are established by distributed networks of sparsely activated neurons, often called engrams (81). Neighboring neurons can have opposing functions or play different roles at various time points (82). Uniform targeting of neurons, even with pathway or cell-type specificity, might therefore not answer ‘*how* the brain works’, but rather how it ‘*can* work’ (83). Therefore, using IEG promotors to selectively express chemogenetic/optogenetic vectors in neurons normally activated during a particular behavior is essential for understanding how endogenous neural activity maps onto the behavioral repertoire of both individual animals, as well as the variability within populations of animals. Is individual variability related to the neurons within a brain region? Or to how neurons are distributed within the whole-brain engram? Alternatively, individual variability could be related to different pathways predominating in different animals. These types of questions can only be answered by activity-dependent selective targeting of neurons. Six of the reviewed studies used a trapping method (cFos tTa, TetTag, TRAP mice) and the use of these technologies is likely to increase overtime, resulting in more refined understanding of neural circuit functions (33,34,84–87).

A prominent example of the role of engram-specificity in circuit dissection is the BLA. While the classical view of the BLA as a “fear” center largely remains uncontested (88,89), activity-dependent labeling has identified subsets of behaviorally activated neurons with distinct roles and functional connectivity (67,77,90,91). Distinct amygdala positive- and negative-valence neurons have been discovered (90). These neurons interact via mutual inhibition and exhibit different functional connectivity. Furthermore, an overlap of positive-valence neurons with fear extinction neurons has been observed (91). Thus, manipulations of the BLA can result in divergent or even opposing behaviors depending on temporally-defined neuronal subpopulations targeted.

Because of the brain’s organization into engrams, conclusions based on pathway or regional manipulations could therefore be either frankly erroneous or fall into the category of ‘what the brain can do’. In a study utilizing activity-dependent tagging to label neurons activated during fear conditioning across 409 brain regions gives weight to this concept. A highly distributed pattern of activity and connectivity was observed, suggesting network redundancy within the brain (87). Single region activation of engram ensembles conferred fear memory recall (albeit not at the same level as multiple engram ensemble activation) demonstrating the ability of neurons within unitary regions to drive behavior despite not acting on their own endogenously (87).

### Same pathway, different action

A visually apparent theme in **Figure 3** is that several pathways promote negative affect with some manipulations and positive affect with others, highlighting the complexity of the brain’s architecture. Increasingly sophisticated targeting specificity have contributed to this pathway duality by improved targeting of small regions, cell-type specificity and, and projection-specificity. This increasing precision has both advantages and disadvantages.

An elegant example of “same pathway, different action” comes from combining projection and cell-type specificity in manipulations of BST→VTA. This pathway contains both glutamatergic and GABAergic projections. Importantly, photoactivation of BST→VTA glutamatergic projections results in aversive and anxiogenic behavior while photoactivation of BST→VTA GABAergic projections produce rewarding and anxiolytic phenotypes (92). However, such precise manipulations can also result in a distorted view, given that the endogenous interplay between BST→VTA GABAergic versus glutamatergic neurons remains poorly understood. Furthermore, this pathway has been implicated in human drug seeking behavior (93), but such studies do not provide data on cellular specificity. GABAergic and glutamatergic neuronal function is intricately interconnected in the brain and therefore, most likely, in the human case both types of neurons are involved. Interestingly, the organization of this pathway has also been found to contain key differences between mice and rats (94) and therefore precise manipulations may not translate between species.

Another cause of discordant results within a pathway comes from increasingly refined subregional localization. As mentioned previously, the mPFC→BLA pathway has been implicated in both fear memory learning and fear extinction learning, with the PL→BLA implicated in the former (9,62) and the IL→BLA responsible for the latter (62,95). Similarly, discrepancies in behavioral outcomes with manipulations of the HPF→Amyg pathway can be attributed to subregional targeting of both input neurons (dentate gyrus (DG) (34) versus the CA1 (32) subregions of the HPF) and output targets (basolateral versus central sub-regions of Amyg (78). Subregional specificity also plays a role in activation of the HPF→mPFC pathway in fear-related behaviors. HPF→IL promotes fear relapse (27), whereas HPF→PL attenuates fear renewal (31). Interestingly, HPF→mPFC activation leads to opposing results in terms of anxiety-versus depressive-like behaviors, potentially indicating affect-specific roles for subsets of neuronal populations in this pathway (28–30).

Additional mechanisms by which the same pathway can display different actions include activity-dependent targeting of neurons for manipulation and the specific timing chosen for the stimulation. As mentioned before, the vHPC →BLA pathway has been implicated in fear memory learning (32, 78). However, the opposite contribution of this pathway was observed when vHPC→BLA neurons were targeted for opsin expression based on their engagement in a positive experience (34).Thus, even neurons of same molecular and projection identity can have opposing contributions to a behavior based on the memory trace they are recruited to. An example of timing effects on behavioral output comes from studies probing the role of the VTA→ACB pathway in social defeat stress. Phasic stimulation during stress and/or the social interaction testing induces susceptibility (35, 36), whereas stimulation after social defeat stress but before the social interaction test induces resilience (37).

Additional mediators of divergent pathway results likely exist. **Tables 3, 4** and MouseCircuits.org provide a simplified way to compare studies for rapid insights into emergent properties of neural circuits. Such insights are vital to the eventual translation of identified rodent circuit function into clinical advances

## GUIDING PRINCIPLES FOR MOVING THE ERA OF AFFECTIVE CIRCUITRY FORWARD

### Methodological considerations

Outcomes of experiments are dependent on a number of methodical choices. For example, the choice of optogenetic versus chemogenetic perturbation can affect conclusions about the role of a circuit. For the most part, the 12 studies (7 pathway focused and 5 region focused) that utilized both optogenetics and chemogenetics came to similar conclusions with both tools. However, subtle differences have also been reported. Chemogenetic activation of vHPC→mPFC during the forced swim test leads to differences in immobility, swimming, and climbing while optogenetic photostimulation results in differences in climbing and swimming only (30). Less subtle differences have been identified in experiments not covered, including differences in specific behavioral outcomes when a circuit is targeted with optogenetics versus chemogenetics (87).

As popularity of activity-dependent labeling grows, it is also important to consider that not all IEGs are made equal. IEGs can have varying patterns of expression across different regions in response to stress (97,98). For example, Covington and colleagues found that optogenetic stimulation increases cFos expression in all conditions tested, but only increases Arc expression following stimulation longer than 30 minutes (97). Others have also found that optogenetic stimulation does not reliably increase Arc expression, potentially due to the complexity of Arc transcription (99,100). Therefore, experimental results following activity-dependent labeling will be partially dependent on IEG choice. Temporal-specific labeling also suffers from significant limitations in trapping window. Most methods have trapping windows of eight to 24 hours, a time-frame which is unlikely to be specific only to the neurons of interest.

It is also important to note that there are technical limitations in manipulations of increasing specificity. Viral spread and infection are tightly coupled to the amount of viral particles injected, which is difficult to precisely control. Viral affinity can differ among both viruses of different serotypes and neuronal types, thereby potentially leading to labeling of nonphysiological ratios of neurons. Labeling based on cell-type specific promoters is also not perfect. For example, CamKII, the choice promoter for selective targeting of excitatory neurons, also leads to expression of virally-packaged proteins in a percentage of inhibitory neurons (64,67).

Taken together, these methodological limitations imply a need for numerous controls to be added to each experimental design. Currently, this is not standard practice. Most commonly, controls involve the expression of a non-opsin/non-DREADD protein to control for injection, viral infection, and photo/drug delivery. An ideal experiment however would include: (1) both chemogenetic and optogenetic manipulation of the region or pathway of interest, (2) bidirectional control to test the pathway under both inhibitory and excitatory conditions, (3) delivery of the opsin/DREADD using viruses of multiple serotypes, (4) systematic dissection of the contribution of each cell-type within the region/pathway of interest as well as all neurons together, (5) “dose-response” analyses to assess the threshold number and location of neurons necessary for driving a behavior using varying amounts of viral particles, (6) “doseresponse” analyses of photostimulation protocol and drug concentration, (7) time-course analyses for “trapping” and/or the delivery of photo/drug stimulation, and (8) investigation of the pathway across multiple behaviors testing behaviors spanning both equivalent and differing affective domains. This type of comprehensive experimental design is imperative for enhanced reproducibility but implausible for an individual lab. A resource such as MouseCircuits.org could therefore aid circuit dissection to move toward this goal as a collaborative open science enterprise by enabling comparisons across studies to effectively generate some of the above mentioned “controls”.

### Sample size considerations

The average number of animals used across all the studies reviewed ranges from 7-13 animals per behavior and manipulation group. In total, the aggregated functional connectome shown in **Figure 3** represents circuit manipulations in 742-1259 animals. In comparison, human studies investigating the role of regions or pathways in behavior or cognition are often based on hundreds to thousands of individuals (101–103). The impetus for the relatively low sample sizes in rodent circuit studies comes from a combination of feasibility of conducting technically demanding experiments with large sample sizes and the universal academic goal of reducing the number of animals used to the minimum. However, if findings from underpowered studies do not replicate, ultimately more resources and animals will need to be allocated to generate the ground truth functional connectome. MouseCircuits.org can help mitigate this risk by generating a centralized and iterative aggregate view of circuit function.

### Sex as a biological variable

Despite the well-documented human preponderance of females afflicted by mood disorders (104,105), the majority of identified studies utilized males only, or did not report the sex of experimental animals. Importantly, out of the 17 studies that included both male and female animals, one study used males for inhibition and females for activation (106), another used females only for tracing studies (107), and a third used males for optogenetic and females for chemogenetic manipulation (108).

On January 25th, 2016, the National Institutes of Health (NIH) implemented a laudable policy requiring investigators to consider sex as a biological variable (SABV) in their grant submissions. SABV was first required for fiscal year 2016 research grant applications, taking effect in fiscal year 2017. Encouragingly, this policy has led to some progress in circuit dissection studies: only 5% (5 studies: 1 regional and 4 pathway) of studies prior to 2017 included females, but 11% (12 studies: 6 regional and 6 pathway) included females after 2017. This trend is likely to improve further and to greatly aid the transnational power of circuit data.

### The need for informed analysis: observation before perturbation

The long-term goal of uncovering the mysteries of affective circuitry is improved understanding of human disorders. To move toward this goal, two guiding principles are necessary in future circuit manipulations: they should be based on identified human functional and structural disfunction, and they should aim to mimic endogenous neural function. The first principle requires a much closer alignment between human and animal research. There is a plethora of human literature on functional and structural connectivity in disordered affect (109–114). These human neural pathways should serve as a blueprint for rodent experimentation, yet most circuit studies to-date base their hypotheses on prior rodent work. As discussed previously, this could lead to propagation of error and movement away from translatability.

The second principle requires thorough examination of normal and abnormal circuit function, followed by careful consideration of experimental conditions that capture identified endogenous activity during perturbation. Is the examined pathway normally activated during the chosen behavior? If so, with what temporal dynamics? Is the signal in this pathway unique to the chosen behavior? Answering these questions involves significant experimental investment prior to chemogenetic/optogenetic manipulations using a combination of *in vivo* electrophysiological or optical recording techniques and *ex vivo* tracing and quantification of activity markers.

Currently, this type of approach is rarely used, with only a minority of reviewed studies reporting observation prior to perturbation of the target region or pathway. Yet poignant examples exist for such prudence moving forward. In particular, the importance of temporal dynamics has been demonstrated in multiple pathways. Stimulation at various time-points within the same behavioral paradigm results in different outcomes in vHPC→Amyg (78), BLA→mPFC (64), VTA→ACB(35,37), and vHPC→NAc (98). An additional example is the inhibition of PL during extinction, which in different studies has either accelerated extinction or had no effect, likely due to timing of manipulation with respect to tone presentation (38,67). Presumably, time-dependence is due to different circuits mediating varying aspects of a behavior. For instance, the PL→BLA pathway is critical for fear retrieval at 6 hours post fear conditioning (115), but at 1 and 28 days, fear retrieval has shifted to the anterior cingulate cortex (ACC)→LA (116).

### Understanding the parts of the sum and the sum of the parts

In an era when both grants and papers are outside of the reach for researchers focused on replicating prior work, we predict the complexity of the single pathway connectome to grow disproportionally to confirmatory studies. This could be particularly troublesome given that the reductionism that has dominated both basic and translational psychiatric research has been increasingly coming into question, with recent evidence that psychiatric diseases might best be interpreted at the level of network emergent properties rather than individual symptoms or the behavioral level (96). There is therefore a crucial need for bidirectional understanding of how the individual components of a circuit contribute to brain-wide activity and how network states influence neuronal function.

The majority of reviewed studies targeted a specific neuronal subpopulation for manipulation of a region or pathway of interest, most commonly glutamatergic neurons expressing CaMKII. Because of the complex interplay between multiple cell-types involved in responding to a stimulus or generating a behavior, individual findings are currently hard to translate to human network function. Behaviors arise from coordinated activity in distributed networks across the entire brain (117). In fear learning for example, it was recently demonstrated that the memory is stored in connected engrams dispersed across the brain (87). While individual circuit findings cannot easily be interpreted in a brain-wide context, as data from individual cell types and pathways is added to a shared resource such as MouseCircuits.org, the interplay is likely to emerge over time.

Similarly, brain-wide connectivity influences the outcomes of manipulations of a specific pathway. Because of complex long-range connectivity, a change in the activity of one region or pathway ripples through the network, shifting activity of other circuits by compensatory or homeostatic mechanisms (118). For example, compensatory pathways can support fear extinction even in the face of a compromised amygdala (119). Network degeneracy, the concept that a circuit generates more than one output and that a pattern of salient neural activity can be generated by more than one circuit (120), also plays a role. Evidence exists for network degeneracy in fear memory circuits. For example blocking dorsal hippocampus (dHPC) via local microinfusion of glutamatergic receptor antagonists disrupts fear memory recall, but the impairment can be overcome by optogenetic activation of a different region – the retrosplenial cortex (121).

The brain-wide effects of stimulation of a particular circuit are hard to predict and very few studies to-date have tackled this question. Using optogenetic stimulation and whole-brain light-sheet microscopy, brain-wide circuit interrogation in zebrafish has shed light on some of these complex interactions (133). Stimulation of one neuronal ensemble was found to increase activity of some brain regions and decrease activity of others. Additionally, inhibiting versus stimulating a particular circuit does not necessarily translate into opposite maps of brain-wide activity changes. Even the time-course of activity changes across the brain can vary for different regions (133). Another recent strategy to tackle this issue is “chemo-connectomics”, which combines functional magnetic resonance imaging (fMRI) with chemogenetics. Using this approach, rapid Resting-State Network (RSN) connectivity changes have been observed following chemogenetic activation of locus coeruleus (LC) (123). These data are imperative in parsing out network effects of chosen circuit manipulations, but difficult to perform and out-of-the reach for most researchers. As data of individual pathways is added to MouseCircuits.org, informed decisions can be made during experimental design, by quickly scanning known upstream and downstream connectivity.

## A SHARED OPEN-SOURCE TOOL FOR MOVING THE CIRCUIT ERA FORWARD

The movement toward open science has generated an abundance of recent resources for the neuroscience community, including the Allan Brain Mouse Connectivity Atlas (122), NeuroMorpho.Org (79, 124–126); GeneNetwork (129), MouseBytes (130), and MouseLight (131). These tools are changing the landscape and culture of neuroscience by maximizing data visibility and impact.

Within this landscape, we envision MouseCircuits.org to aid the translational goal of an integrative view of individual neurocircuit function and whole-brain network organization (132). NeuroMorpho.Org (125) serves as excellent precedent for this vision. As an online repository of neuronal reconstructions from labs around the world, it now hosts over 100,000 neurons from dozens of species and virtually every brain region (79). The number of publications based on secondary analyses of these data currently exceeds the number of original publications for which the neurons were reconstructed (79). Importantly, these secondary analyses have used the raw data of neuronal morphology to generate emergent theories of connectivity in novel ways, e.g. by estimating diffusion tensor imaging (DTI) findings (126, 127). We foresee a repository of functionally dissected individual pathways to lead to emergent properties of other whole-brain imaging modalities, such as fMRI. Ultimately, this will connect rodent data, in which perturbation is possible, to human data, to which we are collectively aiming our clinical advances.

## ACKNOWLEDGEMENTS

This work was funded by 5R01MH111918 to DD and 5T32MH015174 to KA. We thank Allie Lipshutz for her critical assistance with the tables, the entire DOORlab for their support - especially to Maya Erler and Peter Rogu, to the supportive team at Columbia Psychaitry/NYSPI, and to #ScienceTwitter for keeping us up to date on the most recent #circuitera discoveries.

